# Cross-kingdom metagenomic profiling of Lake Hillier reveals pigment-rich polyextremophiles and wide-ranging metabolic adaptations

**DOI:** 10.1101/2022.02.17.480683

**Authors:** Maria A. Sierra, Krista A. Ryon, Braden T. Tierney, Jonathan Foox, Chandrima Bhattacharya, Evan Afshin, Daniel Butler, Stefan J. Green, W. Kelley Thomas, Jordan Ramsdell, Nathan J. Bivens, Ken McGrath, Christopher E. Mason, Scott W. Tighe

**Author notes:** **Corresponding authors:** Christopher E. Mason, Scott W. Tighe.

## Abstract

Background Lake Hillier is a hypersaline lake known for its distinctive bright pink color. The cause of this phenomenon in other hypersaline sites has been attributed to halophiles, *Dunaliella*, and *Salinibacter*, however, a systematic analysis of the microbial communities, their functional features, and the prevalence of pigment-producing-metabolisms has not been previously studied.

Our results are evidence that Lake Hillier is composed of a diverse set of microorganisms including archaea, bacteria, algae, and viruses. Our data indicate a core microbiome in Lake Hillier composed of multiple pigment-producer microbes, many of which are cataloged as polyextremophiles. Additionally, we estimated the diversity of metabolic pathways in the lake and determined that many of these are related to pigment production. We reconstructed complete or partial genomes for 21 discrete bacteria (N = 14) and archaea (N = 7), only 2 of which could be taxonomically annotated to previously observed species.

Our findings provide the first metagenomic study to decipher the source of the pink color of Australia’s Lake Hillier. The study of this pink hypersaline environment is evidence of a microbial consortium of pigment producers, a repertoire of polyextremophiles, a core microbiome and potentially novel species.

## Background

“Extreme” environments are characterized not only by conditions hostile to life (e.g. high temperatures, high salinity) but also often by striking phenotypes, such as color or smell (21; 28; 61). These are often linked to the complex biochemical arsenal required for organisms (i.e. extremophiles) to adapt to antagonistic environments. However, in many cases, the exact sources of these ecosystem-level phenotypes are not known, nor are the taxonomic composition of such environments, the molecular responses of the microbial inhabitants, or the functional adaptations that enable their survival.

Furthermore, in the last four decades, extreme environments have been of great interest for the discovery of new species (10; 14; 86), the study ecological systems (25; 32; 56), evolutionary history (8; 77; 84) and biotechnological purposes (71). The Extreme Microbiome Project (XMP) launched with the goal to develop new methods to detect and characterize novel microbes of different extreme environments (88).

As part of this mission, the XMP has aimed to biologically and biochemically profile Lake Hillier, which is colored bright pink, making it an example of an extreme environment with a readily apparent, yet not fully understood, ecosystem phenotype. Lake Hillier covers an area of 0.15 km^2^ and is located off the coast of Western Australia on Middle Island, the largest of the islands in the Recherche Archipelago Nature Reserve. It is considered “extreme” due to its hypersaline and phosphate-limited nature. It has a salt concentration of 28%, mainly composed of chloride and sodium (Lizamore) and has precipitated salt crystal within the water column. The vast majority of water bodies on Earth are saline, with some lakes and lagoons systems exhibiting this characteristic pink color, however, few reach this salinity level. Oceans contain on average 35 g/l dissolved salts (3.5% salt concentration) (39).

Previous studies in other saline environments have hypothesized similar lake pigments to derive from the algae *Dunaliella salina* (16). *D. salina* is a unicellular microalgae species known for its ability to thrive in high saline conditions and for its production of the carotenoid pigment *β*-carotene. However, there has yet to be a systematic exploration of the microbial communities and their metabolic features that may contribute specifically to Lake Hillier’s pink color in addition to, or perhaps in lieu of, *Dunaliella salina*.

Further, hypersaline environments have been recognized as potential diversity and evolutionary hot-spots (34; 41); they have even been proposed as Martian analogs due to their similar chemical composition to Mars (58). Other studies have characterized the microbial communities of other hypersaline lakes in Australia, revealing novel ecology systems, unique adaptations and rich microbial community structures (26; 27; 35; 69; 70). These studies have shown that one of the most abundant species belong to the genus *Salinibacter*, but other novel microbial species have been also recovered.

Therefore, exploring the biology of Lake Hillier can address several goals beyond identifying the cause of its color: it may serve as a wellspring of potential novel biochemical features and functional elements that enable survival in hypersaline environments. Here, through a shotgun whole genome sequencing-based and metagenomic assembly approach, we (the XMP team) aimed to characterize the microbiome of Lake Hillier in order to partially explain its coloration and also measure its metabolic potential as a function of the algae, bacteria, archaea, and viruses that inhabit it.

## Materials and Methods

### Sampling and environmental parameters

Water and sediment samples were collected in February 2015 from Lake Hillier. The lake is located on Middle Island (34.0950° S, 123.2028° E), the largest of the island in the Recherche Archipelago Nature Reserve, off the southern coast of Western Australia. It is a terminal, perennial lake measuring 600 m in length and 250 m in width. The lake itself is fed by a combination of fresh and brackish groundwater with minimal overland flow due to frequent rainfall. The lake is most notably hypersaline and is permanently pink in coloration but often experiences varying intensities of color depending on seasonal influence.

Samples were collected in the summer, during a time when the lake levels were reduced due to evaporation, leaving a salt-crusted shoreline. Sampling of the bank sediment was carried out on the northern shoreline (34.093330° S, 123.200965° E) and the lake sediment, top (surface) water, and mid-depth water were collected at the same point at the center of the lake (34.093985° S, 123.200966° E). Water temperature and pH were simultaneously measured using the OW-618 indicator (OWAY, PH-618) measuring the temperature to be 26°C and pH to be 7.4. At each sampling site, samples were collected in triplicate in a sterile 1000 mL Duran laboratory bottle (Duran laboratory bottle, Cat.: Z305200). All water was collected by immersion below the surface body of water until filled, while the sediment was collected by manually scooping the upper 3-5 cm layer and draining off all residual lake water that may have been captured before sealing. All sample collection was performed using DNA-free sampling techniques. Water and sediment samples separately went through a filtration process and passed through a sterile membrane filter with a pore size of 0.2 μm (Advantec Membrane Filter, Cat.: A020H047A). Preservation of the retained specimens was divided into three separate methods, mixed into a concentration of ethanol (40%), a Dimethyl Sulfoxide (DMSO) solution, or snap-frozen over dry ice. These preserved samples were stored in 15 mL conical tubes, where they were transported and stored at −20°C until downstream analysis.

### Samples processing and sequencing

#### DNA extraction and sequencing

Due to the hypersaline nature of Lake Hillier, the water samples contained excessively high levels of semi-precipitated sodium chloride crystals. The samples with crystals were dissolved in 5 volumes of molecular grade water to dissolve all precipitated salt and then filtered through a hydrophilic polycarbonate membrane with a pore size of 0.2 μm (Millipore Sigma, Cat.: GTTP04700) using a glass filtration apparatus. The glass apparatus was baked and sterilized prior to use to eliminate trace levels of DNA. The filter membranes were placed in a 50 mL conical tube with 5.0 mL of 1X Phosphate-buffered saline (PBS) (Cytiva, Cat.: SH30910.03) and mechanically homogenized using a bead beating grinder for 1 min at 4000 reps (MP Biomedical, FastPrep24). After homogenization, the sample was transferred to (3) 1.5ml microcentrifuge tubes (Axygen, Cat.: MCT-175-C) and centrifuged at 1500 x g for 5 min to pellet. The supernatant was removed and the pellet was resuspended in 50 μL of PBS. The sediment samples, approximately 200 mg, were washed 5 times in 1.5 mL of molecular grade water (to reduce the saline content) and pelleted by centrifugation at 1500 x g. After centrifugation, the supernatant was removed and the pellets were resuspended in 500 μL of 1X PBS. The water and sediment samples, 100 μL aliquots, were then heated for 10 min at 80C in order to inactivate the DNase activity. After heating, 20 μL of a multilytic enzyme mix (Millipore Sigma MetaPolyzyme, Cat.: MAC4L) and 5 μL of (2%) sodium azide was added to each sample. The samples were incubated at 35°C for 12 hrs to digest microbial cell walls. DNA was extracted from the resulting digests using the E.Z.N.A Mollusc DNA Kit (Omega Bio-tek, D3373-01), following manufacturers instructions.

DNA sequencing libraries were prepared using a Nextera XT library kit (Illumina, Cat.: FC-131-1024) with 1 ng of DNA input. A total of two DNA samples of sediment and water samples were subjected to metagenome shotgun sequencing. Sequencing was done at the Hubbard Center for Genome Studies, at the University of New Hampshire and the University of Vermont using the Illumina HiSeq 1500/2500 DNA sequencer with single-end 100bp reads and the MiSeq System with paired-end 250bp reads.

#### Amplicon Sequencing

For 16SrRNA and 18SrRNA amplicon profiling, DNA was extracted using the PowerSoil DNA Isolation Kit (Qiagen/MO BIO Laboratories, Inc., Cat.: 12888), according to the manufacturer’s protocols, after sample pre-processing described above. To characterize the bacterial and archaeal communities, the small-subunit (SSU) region of the 16S ribosomal DNA (rDNA) gene was amplified using primers broadly targeting bacteria and archaea: 341F (5’-CCTAYGGGRBGCASCAG-3’) and 806R (5’-GGACTAC-NNGGGTATCTAAT-3’) modified on the 5’ end to contain the Illumina Nextera Adaptor i5 & i7 Sequences. PCR reactions were performed using AmpliTaq Gold Master Mix (Applied Biosystems, Cat.:4398881) according to the manufacturer’s recommendations. PCR amplification was performed using the following cycling conditions: 95°C for 5 min; 29 cycles of 94°C for 30 s, 50°C for 60 s, 72°C for 60 s; with a final extension of 7 min at 72°C. The resulting PCR amplicons were purified with Agencourt AMPure XP Beads (Beckman Coulter, Cat.: A63880). A second PCR step was performed with Illumina Nextera XT Index Kit v2 (Illumina, Cat.:FC-131-2001) and Ex Taq DNA Polymerase (TaKaRa Bio, Cat.:RR001), to add sequencing indexes to each amplicon. The final amplicons were cleaned with Agencourt AMPure XP Beads and quantified using PicoGreen (Invitrogen). A total of 48 samples were pooled and prepared by mixing the amplicons in relative concentrations following DNA quantification to prevent ambiguity in the expected coverage. Pooled samples were quantified using the KAPA Biosystems qPCR library quantification Kit and normalized to 4 nM prior to sequencing. Sequencing was performed at the Australian Genome Research Facility (AGRF), University of Queensland, using the Illumina MiSeq System and MiSeq Reagent Kit v3 with paired-end 300bp reads.

Preparation of the archaeal 16SrRNA enriched libraries employed two rounds of PCR. This method was preferential to the amplification and sequencing of a 16S rRNA region specific to archaeal targets. The small-subunit rDNA was first amplified by PCR using a forward primer based on the A2/519R primer sets; positions 2-21 (5’-5’-TTCCGGTTGATCCYGCCGGA-3’) and the reverse primer corresponding to the complement position of 1510-1492 (5”-GGTTACCTTGTTACGACTT-3’) (73). PCR amplification was performed using the following conditions: 95°C for 10 min; 35 cycles of 95°C for 30 s, 60°C for 15 s, 72°C for 50 s; with a final extension of 5 min at 72°C. In total, 16 samples were pooled and sequenced which included 10 sediment and 6 water samples. The second round of PCR followed the manufacturer’s guidelines for 16S rRNA Metagenomic Sequencing Library protocol (Illumina, Cat.: 150442223) for the MiSeq system. The V9 region of microbial eukaryotes 18S rRNA gene was amplified with primer constructs containing universal primers 1391f (5’-GTACACACCGCCCGTC-3’) and EukBr (5’-TGATCCTTCTGCAGGTTCACCTAC-3’). DNA was amplified and the resulting PCR reaction was quantified using the PicoGreen (Invitrogen). The resulting libraries were then individually normalized to 4nM and pooled prior to sequencing. Sequencing was performed at the Australian Genome Research Facility (AGRF), University of Queensland, using an Illumina MiSeq System using the MiSeq Reagent Kit v3 with paired-end 300bp reads.

#### Microscopy and taxonomic classification of cultures

Water and sediment samples were microscopically evaluated using standard wet mount bright field microscopy using 100, 200, and 600X magnification (Ziess AxioPlan 2, Jena, German) to observe algae and other notable biologicals. Image capture was performed using standard photomicroscopy. Culturing of sediment and water samples was performed using a non-quantitative spread and streak plate methods on two types of media types, Marine Broth Agar 2216 (MBA) (BD Difco, Cat.: DF0791-17-4) and Marine Broth Agar 2216 supplemented with 10% NaCl and water recovered from Lake Hillier. All media were inoculated with 20 μL, 30 μL, and 100 μL of water and wet sediment samples and incubated at 22°C and 28°C for 2 weeks until colonies were visually identified. The resulting colonies were photographed (Figure S8) and all isolates were subcultured on MBA 2216 with 10% NaCl. DNA was extracted from pure colonies using the E.Z.N.A Mollusc DNA Kit (Omega Bio-tek, D3373-01) by transferring a loopful of cell mass to 100 μl of Phosphate buffered saline (PBS) buffer and pre-digested with Metapolyzyme (Millipore Sigma, Cat.: MAC4L) for 4 hours prior to bead beating and column extraction. Extracted DNA was quantified with the Qubit spectrofluorometer (ThermoFisher Scientific) and checked for quality using a NanoDrop spectrophotometer to determine protein (260:280 ratio) and salt contamination (230:260 ratio).

Taxonomic identification of isolates was accomplished by PCR amplification of full-length 16srDNA using primers 27F (5’-AGAGTTTGATYMTGGCTCAG-3’) and 1492r (5’-GGYTACCTTGTTACGACTT-3’), digesting withExoSAP-IT (ThermoFisher Scientific, Cat.:78201), followed by Sanger sequencing. Sequencing was performed at the University of Missouri and University of Vermont DNA core facility using an ABI 3730XL Genetic Analyzer (ThermoFisher) (82). Resulting sequences were pairwise-aligned against the 16S_ribosomal_RNA database (with -max_target_seqs=10), and query and subject sequences were aligned with MUSCLE (24) using AliView (49). A phylogenetic tree was inferenced by maximum likelihood (GTR+G4+F -bb 1000) using IQ-TREE (60). The resulting Newick tree was plotted with ggtree (99) (See data availability).

### Bioinformatic Analysis

#### Amplicon reads processing

Paired-ended reads from the 48 samples and all primers for 16S and 18S were quality checked using FASTQC (1). For reads from primers, 27F-519R reverse reads failed quality scores, therefore only forward reads were kept. Both forward and reverse reads from primers 341F-806R and 1391f-EukBr were kept. The remaining reads were processed with the Quantitative Insights into Microbial Ecology Version 2 (QIIME2 v.2020.2) (13). Each primer set was processed independently and then merged into a single set. Briefly: Reads were imported with as –type ‘SampleData[PairedEndSequencesWithQuality]’ –inputformat CasavaOneEightSingleLanePerSampleDirFmt for paired-ended reads, and ‘SampleData[SequencesWithQuality]’ for single-ended reads. Depending on quality scores, reads from each primer set were trimmed independently using qiime dada2 denoise-paired and denoise-single. For reads from primer 27F-519R –p-trunc-len 280 –p-trim-left 20 was used. While –p-trunc-len-f 280 –p-trim-left-f 15 –p-trim-left-r 15 –p-trunc-len-r 210 was used for primer 341F-806R and –p-trim-left-f 20 –p-trim-left-r 30 for 1391f-EukBr.

For taxonomic classification, a classifier from SILVA v1.38 database compatible with QIIME2 was built using the plugin RESCRIPt (74). The classifier was trained with specific region-primers using qiime feature-classifier extract-reads with -p-f-primer AGAGTTTGATCATGGCTCAG –p-r-primer GGACTACHVGGGTWTCTAAT and qiime rescript dereplicate –p-rank-handles ‘silva’ –p-mode ‘uniq’. The classifier was tested using feature-classifier classify-sklearn and biom and taxonomy tables were generated. Further analyses of abundance and diversity were performed in R and Python by custom scripts (See data availability).

#### Whole genome reads processing

Quality controlled was performed on the raw reads from two shotgun-sequenced samples (from sediment and water) via the following steps: BBMap (15) was used to deduplicate and clump reads, and BBDuk was used (within the BBMap suite) to remove adapter contamination (clumpify: optical=f, dupe-subs=2,dedupe=t, bbduk: qout=33 trd=t hdist=1 k=27 ktrim=“r” mink=8 overwrite=true trimq=10 qtrim=‘rl’ threads=10 minlength=51 maxns=-1 minbasefrequency=0.05 ecco=f). Finally, BBMap’s tadpole was applied to correct sequencing errors (mode=correct, ecc=t, ecco=t).

A combination of assembly and short reading mapping approaches were used for the analysis. Kraken2-build was used to construct a custom Kraken2 database (94). This contained the National Center for Biotechnology Information’s (NCBI’s) bacterial and viral RefSeq databases as well as the complete set of GenBank’s Protozoan genomes, the GenBank algal plant genomes, and all fungal genomes. Kraken2 default settings were selected to compute the presence of different taxonomic species in two quality-controlled samples (Sediment and Water), and then Bracken2 (53) to estimate the abundance of these organisms, the database built and software was run with the default settings. To compute pathway abundances, HUMAnN 3.0 (9) was run with default settings. For assembly-based analysis, *de novo* assembly of quality-controlled reads into contigs was performed using metaSPAdes (62) with default parameters, and assembly quality was checked with MetaQUAST (57) (Supplementary Table 1).

#### Extremophile profiling

A list of species present in all sample types (bank, water, and sediment) and sequencing methods (amplicon and WGS) was generated. A cladogram of these species was built using the Environment for Tree Exploration (ETE) toolkit (40) and extremophile profile was incorporated using ggtree (98). To generate the extremophiles profile, the latest version (unpublished) of our database, The Microbe Directory (TMD) was used (78). The Microbe Directory is a database that contains morphological and ecologic characteristics of microbes and is based on published literature and manual curation (54). Using the list of extremophiles from TMD, species found in Lake Hillier were classified into eleven extremophile types: Acidophile, Thermophile, Alkaliphile, Halophile, Psychrophile, Metallotolerant, Oligotroph, Radioresistant, Barophile, Hypolith and Xerophile.

#### Binning and characterizing Metagenome-Assembled Genomes (MAGs)

Metagenome-Assembled Genomes (MAGs) were constructed from assembled contigs in the two wholegenome-sequenced samples using an ensemble binning approach. MetaWRAP (89) was used with default parameters to generate genome bins from CONCOCT (3), MetaBAT (45), and MaxBin2 (95). dRep (64) was used with the following settings: -comp 50 -pa 0.9 -sa 0.95 -nc 0.30 -cm larger, which wraps CheckM (67), to filter these bins (by default removing genomes with >25% contamination) and collapse those remaining into the reported set of non-redundant set of 21 genomes, with genomes sharing greater than 95% Average Nucleotide Identity (ANI) being considered the same taxon. We reported all genomes with completeness >50%, defining genome quality based on the literature (93), with medium quality being between 50% and 90% completeness (and <5% contamination) and high quality being >90% completeness (and also <5% contamination). Taxonomic classification of the resultant low, medium, and high quality MAGs with completeness >50% was done using GTDBTk’s classification workflow running the default settings (17), assigning them best possible taxonomies and placing them in the Genome Taxonomy Database’s bacterial and archaeal trees using ggtree (98).

#### Genome Mining of Metagenomes

The standalone version of AntiSMASH5 v5.2.0 (12) was used to identify Biosynthetic Gene Clusters (BGCs) from the metagenomes assembled by metaSPAdes with the following parameters: –cb-general –asf –smcog-trees –cb-knownclusters –cb-subclusters –pfam2go –taxon bacteria. Using Prodigal (42) for bacterial gene prediction, AntiSMASH was used to predict the BGCs and define them within chemical classes (henceforth, referred to as class). Reports of similarity with any other known BGC based on the MIBiG2 database (46) were also generated. As MIBiG2 database contains all annotated and known BGCs, novel BGCs were defined with less than 80% sequence similarity to MIBiG2 sequences (47). Big-SCAPE/CORASON (59) was used to explore the diversity of BGCs classes predicted by AntiSMASH (parameter: “–mibig”). This grouped the identified BGCs based on similarity networks of gene cluster family (GCF) according to protein family (i.e. Pfam directory). GCFs (referred to also as families) encode for similar secondary metabolites.

## Results

### Diversity of Lake Hillier includes microbes from four domains

A total of 48 samples were collected from sediment and water using three fixation methods (ETOH, FRESH, DMSO), Table **??**. Using two sequencing techniques, A total of 4,563,633 and 186,016,408 sequence reads in amplicon and WGS were obtained, respectively. Sequences were classified into four domains: Archaea, Bacteria, Eukaryota, and Viruses, Figure 1A. Since both sequencing methods differ in scale, the abundance of taxa was log-normalized for both methods independently before comparison. These four microbial groups were similarly abundant in all sample types, however, given the sequencing approach, the presence or abundance of phyla was influenced. For example, bacteria phyla such as Zixibacteria, Sumerlaeota, Patescibacteria, Modulibacteria, Acetothermia and Hydrogenedentes were only found by amplicon (Supplementary Figure S2), while Thermodesulfobacteria, Kiritimatiellaeota, Dictyoglomi, Caldiserica, Aquificae and Bipolaricaulota were only found by WGS.

**Figure 1:**
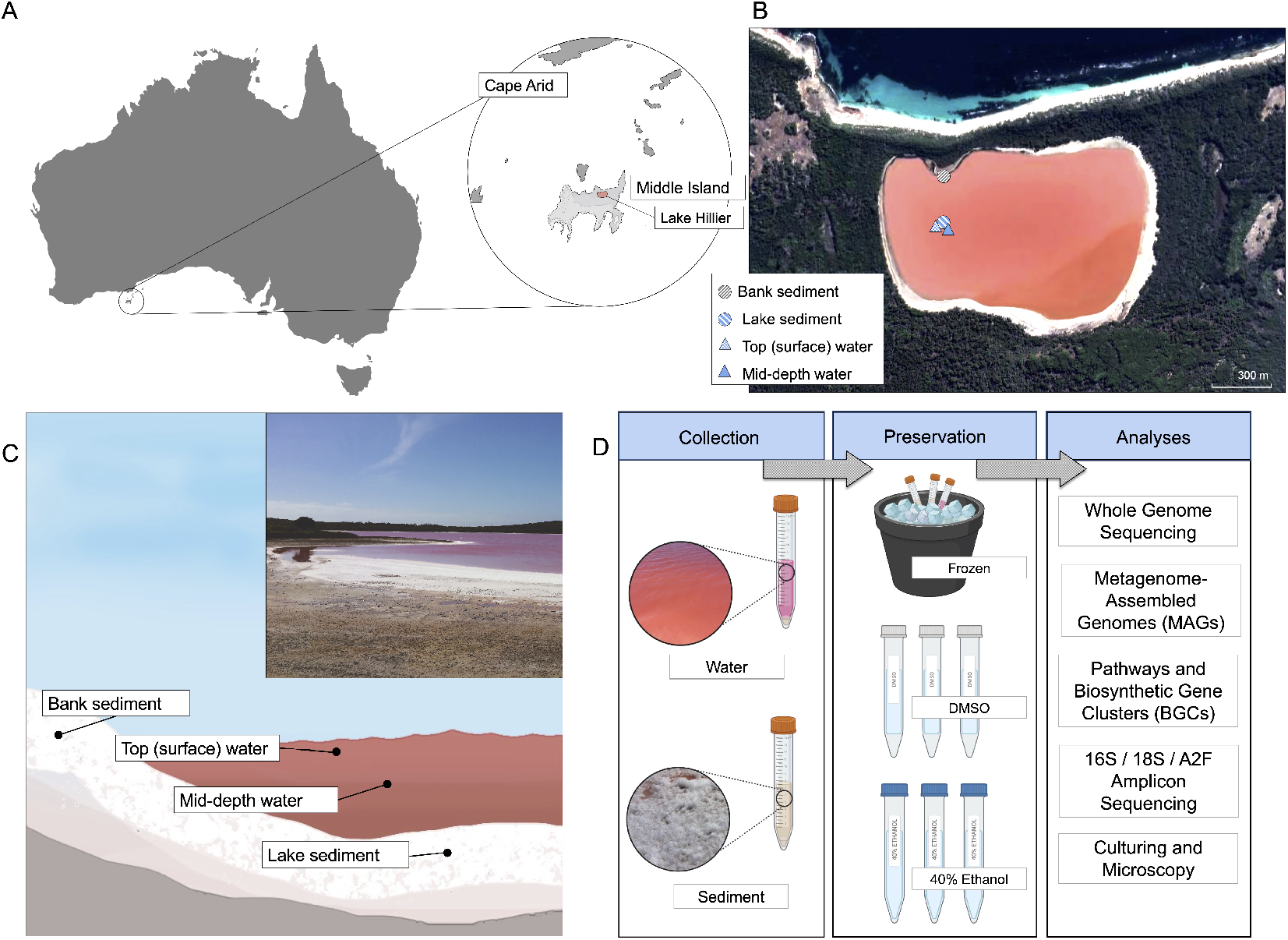
A.Map of Australia. The area circled is part of the Recherche Archipelago (Bay of Isles) off the southern coast of Western Australia, which contains Middle Island and the study site of Lake Hillier.B.Map of the study area. Each icon shows the approximate location of the sampling sites where sediment (bank and lake) and water from the top (surface) and mid-depth was collected.C. Diagram showing an illustrated cross-section of Lake Hillier and an image from the day of sample collection. Each dot shows the approximate location where each sampling type was collected.D.An overview of the sample collection and preservation workflow, where aliquots of the water and sediment samples were preserved in EtOH (40%), DMSO, or frozen over dry ice following collection. The samples were then processed and analyzed using shotgun metagenomics, 16S and 18S rRNA gene sequencing, and archaeal 16S (A2F) rRNA amplification.

Similarly, the prevalence of some phyla in Archaea varied by the sequencing method, (Supplementary Figure S3A). Candidate phyla Hadarchaeum, Altiarchaeota, Aenigmarchaeota and Asgardarchaeota were only found through amplicon sequencing. In contrast to Lokiarchaeota, Korarchaeota, and Thaumarchaeota that were only found by WGS. Eukaryotic members of Labyrinthylomycetes, Apicomplexa, Aphelidae, Ancyromonadida and Amoebozoa were only found by amplicon, while Haptista, Foraminifera and Evosea only by WGS.

Although viruses were only detected through WGS, the presence of 12 phyla was observed (Supplementary Figure S3C). Phylum Uroviricota was the most abundant in both sediment and water. Phyla Teleaviricota, Saleviricota, Duplornaviricota and Artverviricota were only present sediment, while Kitrinoviricota and Artverviricota only in water.

While most phyla showed similar abundances by both sequencing methods, this changed at the species level, (Supplementary Figure S4). The number of unique and shared species varied among domains, Figure 2B. Amplicon and WGS only shared 6 species of Archaea, 51 of Bacteria, and 7 of Eukaryotes. Despite the low number of shared species, most abundances were similar regardless of the sequencing method. All 6 shared members of archaea belonged to the class Halobacteria, and these shared taxa were among the 20 most abundant total species. In bacteria, *Salinibacter ruber* was the most abundant species, followed by members of phylum Proteobacteria. The most abundant Eurkaryote in both methods was the algae *Dunaliella salina*.

**Figure 2:**
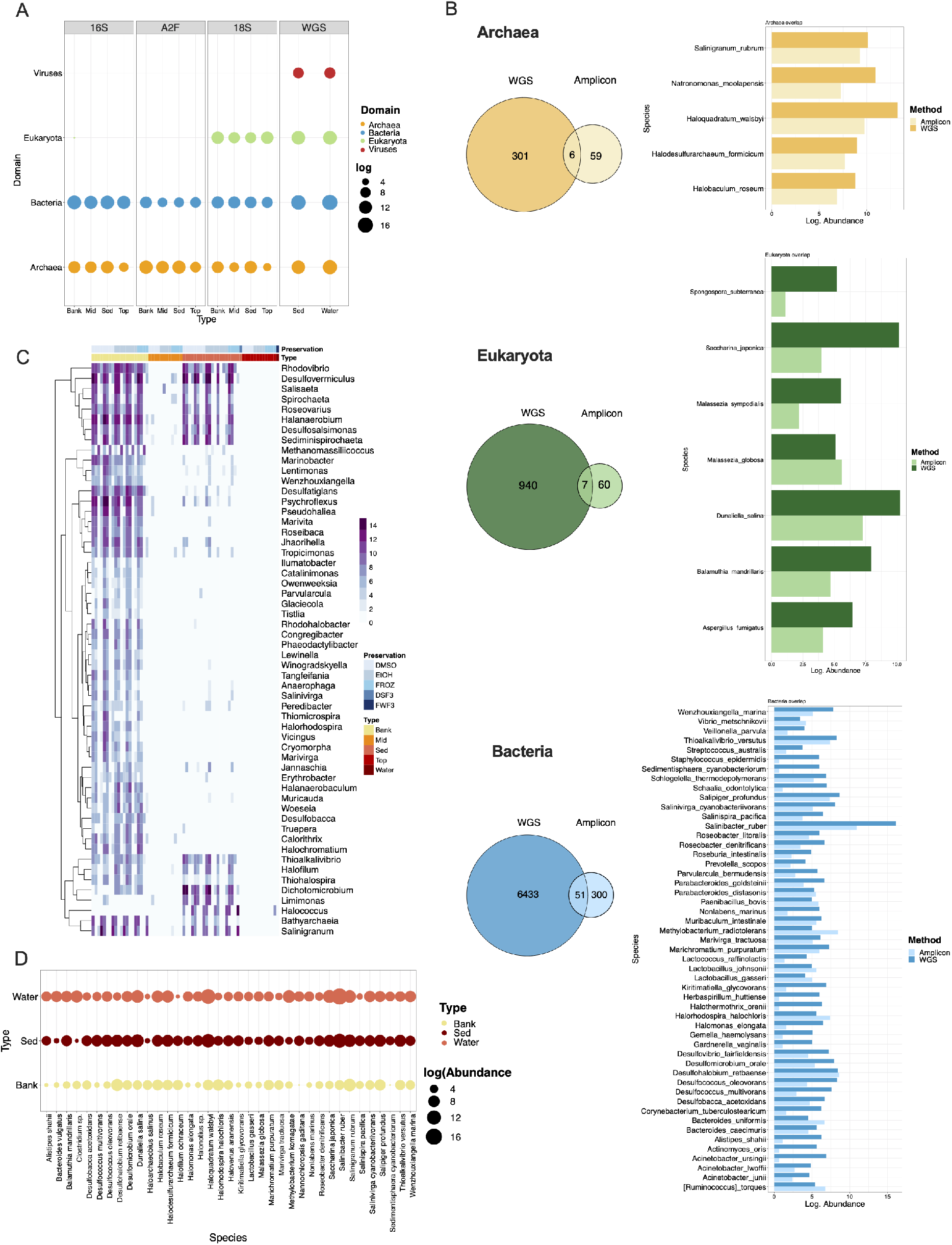
A. Abundance of four domains found by different molecular and sequencing approaches. Abundance log transformed. B. Unique and overlapped species found in Archaea (Top) and Bacteria (Bottom) by both methods (Amplicon and WGS). Taxa found in both methods is represented on the right of each Venn diagram. C. 55 genera with significantly differential abundance among sample types by ALDEx2. Only taxa classified at genus level were included. D. “Core” species: Abundance of taxa found in all sample types.

These results also suggest that the abundance and prevalence of genera do not depend on the preservation method. However, we found significant differential abundances in 124 genera using ALDEx2 (30), based on the sample type and origin, Figure 2C. Samples collected in the bank had most of the 124 genera detected by ALDEx2 and in higher abundance compared with other sample types. Notably, none of these genera were observed in the water samples.

Since Lake Hillier represents a unique halophilic biome, and according to our results, some microbes display a preference for sediments or water, the question of whether there is a set of microbes consistently present in the lake remained. From the 13,909 microbial species found in the lake, 53 species were shared among water, bank, and sediment samples (Supplementary Figure S5). However, only 37 of these species were classified, Figure 1D, belonging mostly to archaea.

### Lake Hillier a source of polyextremophiles

The 498 extremophiles profiled by the Microbe Directory database showed that the most abundant types in Lake Hillier were halophiles with 249 species and thermophiles with 175. Additionally, these results identified the presence of 63 Acidophiles, 49 Psychrophiles, 43 Alkaliphiles, 11 Barophiles, 7 Xerophiles, 7 Metallotolerants, 7 Radioresistant, and 1 Hypolith species.

Some species were profiled to multiple types of extremophiles, such as the case of the thermohalophile *Ignicoccus islandicus*, thermohaloalkaliphile *Natranaerobius thermophilus*, thermoacidophile *Sulfolobus acidocaldarius*, acidophile-metallotolerant *Rhodanobacter denitrificans*, metalloradiationresistant *Herminiimonas arsenitoxidans*, among others. According to TMD, these extremophiles have been found in a range of different types of microbiomes from urban environments, soil, water, to ocean depths, geothermal hot-springs, deserts, and polar environments.

Also, the presence of multiple sulfur-reducing bacteria from the order Chromatiales was observed. This order is also known as purple sulfur bacteria (PSB). To further characterize these species, we downloaded the complete list of species of PSB as well as purple non-sulfur bacteria (PNSB, family Rhodospirillaceae) from NCBI Taxonomy and matched it with our a full list of species. These comparisons resulted in 55 PSB species and 15 PNSB being identified in the Lake Hillier data, Figure S6. From the PSB species found, 9 taxa were profiled as at least one type of extremophiles: Halophiles *Aquisalimonas halophila*, *Nitrosococcus halophilus*, *Spiribacter curvatus*, *Spiribacter salinus*, *Thiohalobacter thiocyanaticus*, *Thiohalospira halophila*, thermohalophile *Halorhodospira halochloris*, alkalohalophile *Thioalkalivibrio sulfidiphilus*, and thermophile *Thermochromatium tepidum*, and some were also members of the core microbiome (Figure 2D).

### Lake Hillier as a source of novel and diverse Metagenome-Assembled Genomes

WGS shotgun data were processed for high-quality Metagenome-Assembled Genomes (MAGs). These MAGs were grouped into 3 categories: high quality (completeness >90 % and contamination <5 %), medium quality (completeness >50 % and <90 %), and low quality (contamination >5 % and <25 % and completeness >50 %). After removing redundant genomes that were at least 95 % in terms of Average-Nucleotide-Identity (ANI), 1 high quality genome, 9 medium quality genomes, and 11 low quality genomes were identified, Figure 4A, Supplementary Table 1.

**Figure 3:**
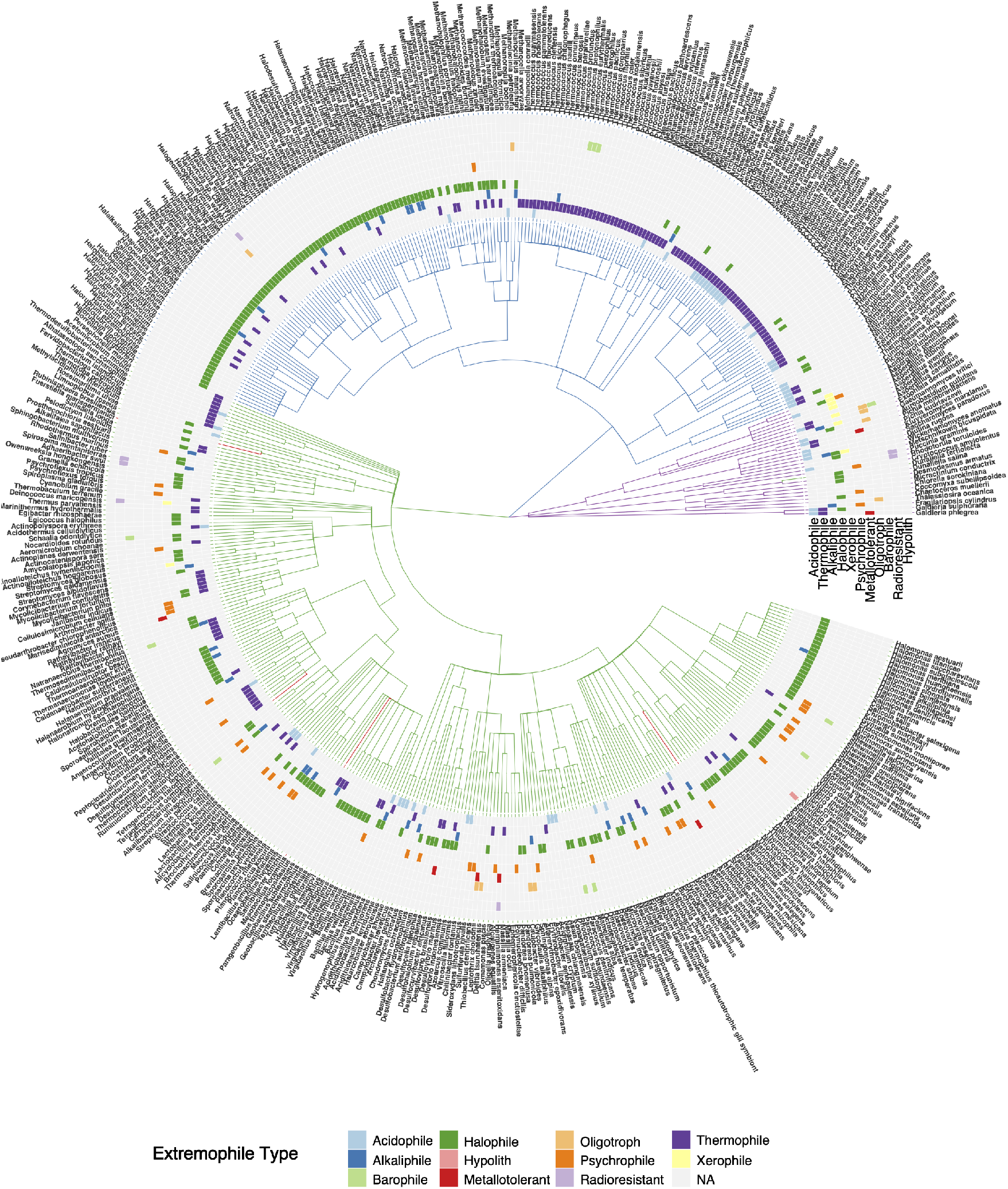
Tree of extremophiles: 472 species were found in the Microbe Directory database to be at least one type of extremophile. Branch color represent domain to which the microorganism corresponds. Heatmap represents the types of extremophile associated to each taxa.

**Figure 4:**
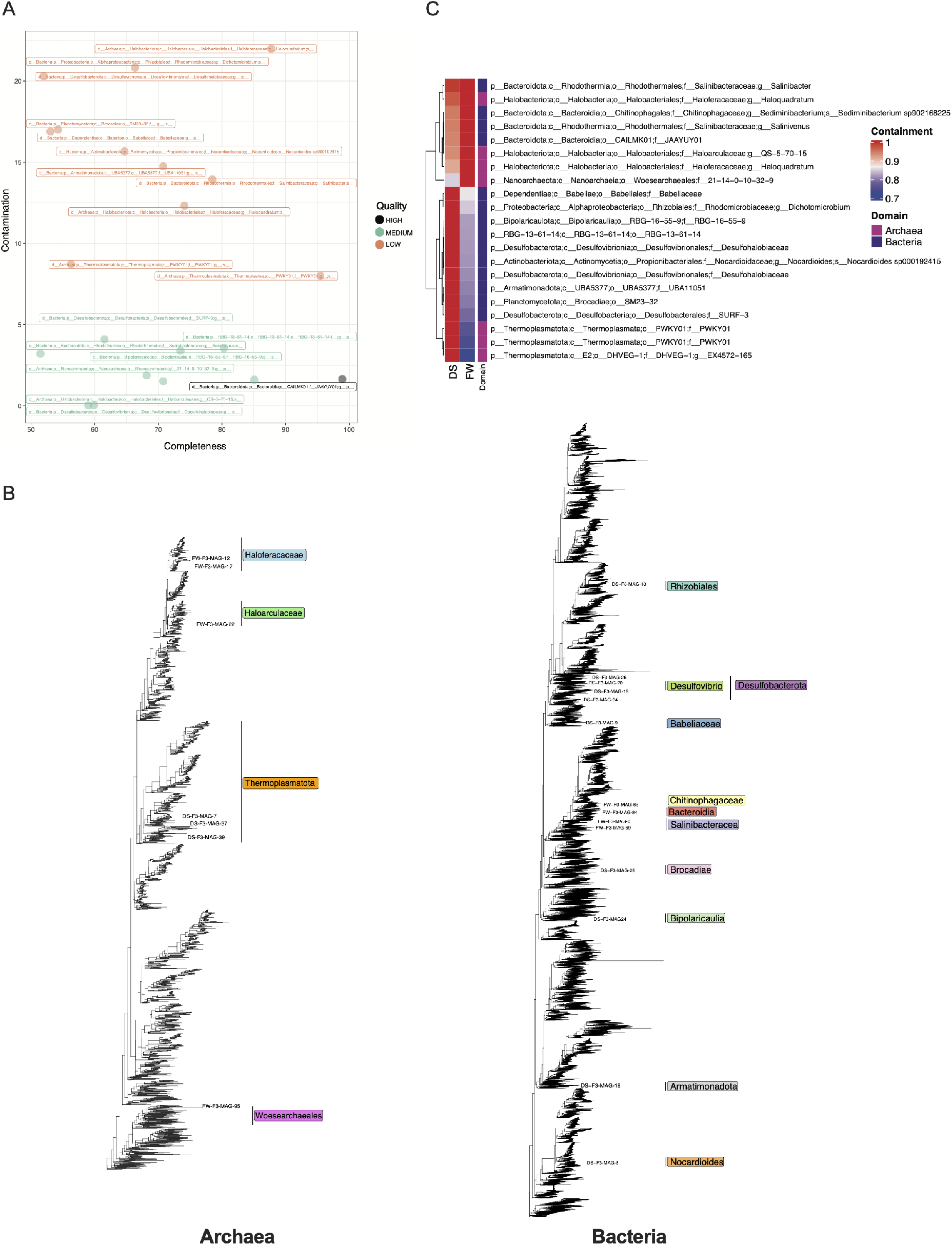
A. MAG quality. MAG completeness and contamination, as reported by dRep, are indicated on the X and Y axes. Each point represents 1 of 21 non-redundant MAGs. These are colored by our definitions of high (>90% completeness and <5% contamination), medium (between 50% and 90% completeness and <5% contamination), and low (between 50% and 90% completeness and >5% contamination) quality. B. Trees of MAGs position on the Archaea (left) and Bacteria (right) phylogeny. C. Containment values within the two WGS samples (determined using Mash screen) of the 21 non-redundant MAGs colored by assigned domain.

These 21 genomes were annotated according to the Genome Taxonomy Database (GTDB) and only two MAGs were ascribed to species level taxonomy: (*Sediminibacterium sp902168225* and *Nocardioides sp000192415*), with the former being medium quality and the latter being low quality. Seven potential MAGs were annotated as archaeal and 14 were annotated as bacterial in origin, Figure 4B. The high quality genome was annotated as a member of the *JAAYUY01* family (within the Bacteroidota phylum). In addition to *Nocardioides sp000192415*, which is similar to the organism with the dominant uniquely abundant functions according to the pathway analysis, we assembled MAGs for a potential member of the genus *Salinibacter*, which corresponded to the functionally dominant water genus, Supplemental Figure S7.

Containment analysis was accomplished using Mash screen (65) to identify which samples likely contained our 21 bins based on the frequency of metagenome-genome kmer matching. We identified similar results to our short-read and amplicon analysis, with the sediment and water samples containing distinct, but overlapping, bin representation, Figure 4C. The sediment samples contained matches to nearly all bins, whereas the water samples had fewer, but higher confidence (i.e. likely higher abundance) matches. The *Salinibacter* genus and *Nocardioides sp000192415* were uniquely present in the water and soil samples, respectively. Archaeal and bacterial genomes were split equally between the two samples, though the clades were distinct such that all three Thermoplasmatota representatives were in the soil sample, whereas the water sample was dominated by Bacteroidota and Halobacteriota.

### Estimating the metabolic capacity of Lake Hillier

The abundance of metabolic pathways in the two WGS samples from water and sediment were estimated and found to contain variable metabolic potential ascribed to distinct species, Figure 5A. In this analysis pathways were split into three groups, 1) mutually abundant in both samples, 2) abundant in sediment but not in water, and 3) abundant in water but not in sediment. Overall, the sediment had a higher abundance of pathways that were not found in the water metagenome. These pathways were almost universally dominated by an unclassified Nocardioidaceae species, a member of the order Corynebacteriales. Mutually abundant pathways tended to be core microbial metabolism that could not be assigned to a specific species. This included, for example, pathways related to nucleotide biosynthesis.

**Figure 5:**
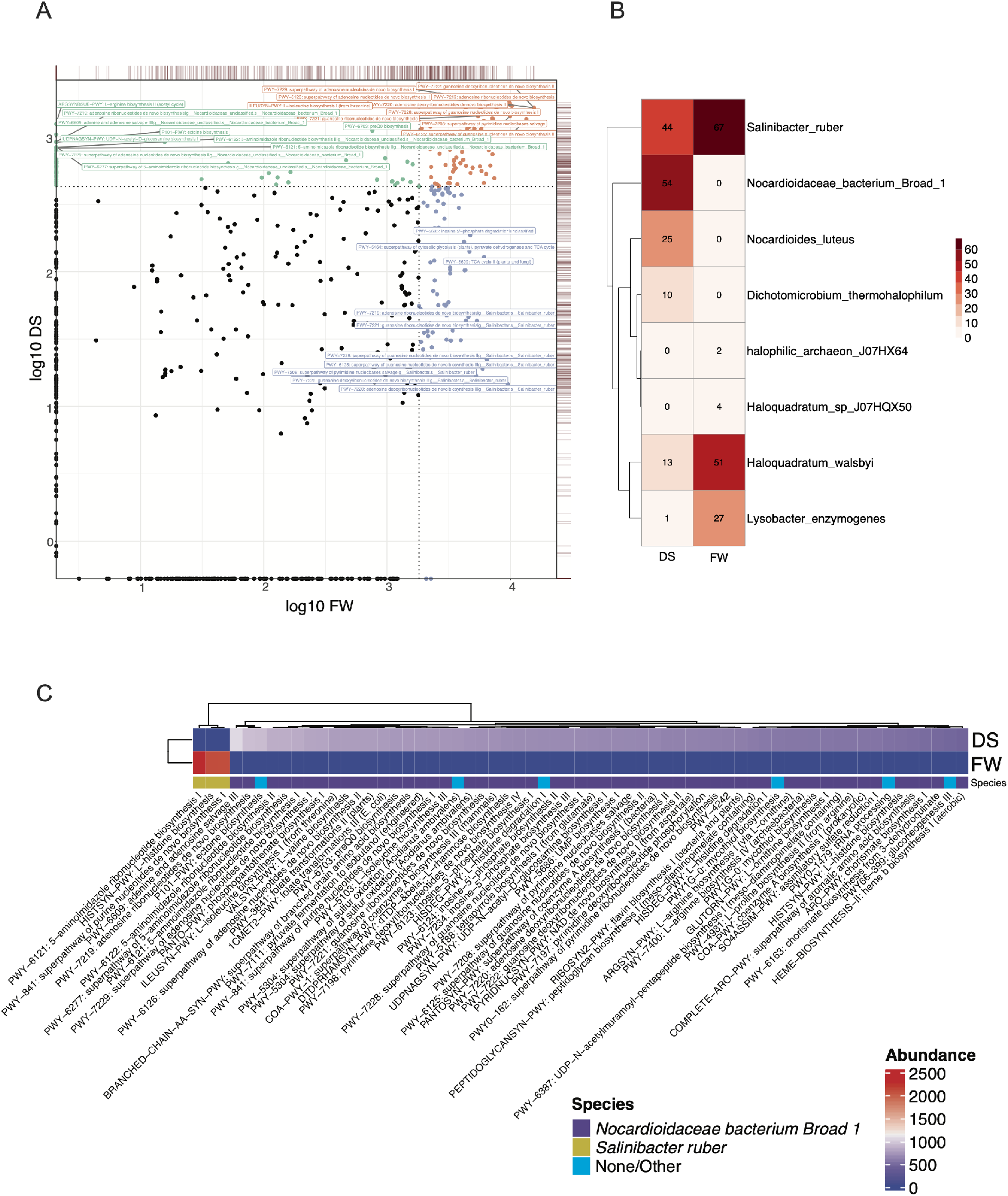
A. Pathway log-abundance of the two metagenomes from water (X axis) and sediment (Y axis), colored by most abundant pathway in each sample: Green sediment, purple water, and orange moth abundant pathways in both samples. Dotted lines indicate the 0.75 quantile of log-abundance for the X and Y axes. Red lines on top and right hand side of plot indicate the density of points for the corresponding axes. B. frequency of the different pathways assigned to microbial species by HUMAnN 3.0. C. Pathway abundance in sediment (DS) and water (FW) in two of the most reported species by HUMAnN 3.0.

These analyses additionally identified pathways assigned to species arising predominantly in the WGS samples, Figure 5B. Once more, *Salinibacter ruber* and the unclassified Nocardioidaceae taxon were among the two species with the highest number of distinct pathways. Functionally, there were a number of uniquely high abundant and diverse pathways annotated to these taxa (Figure 5C), including ectoine biosynthesis, heme biosynthesis, chorismate biosynthesis, pyruvate fermentation to isobutanol, and various folate transformations. Additionally, a number of pathways related to sulfur oxidation were observed, which relates to the previous observation of purple sulfur bacteria.

The water sample was notably distinct in the sense that it was dominated by *Salinibacter* associated pathways. There were fewer (3) pathways that were in the top quantile of abundance for this sample but not found at all in the sediment sample. However, these three all annotated as *Salinibacter* core functions, were in much higher abundance than any of the uniquely abundant pathways in the sediment (Figure 5C). Other abundant, non-core pathways annotated to *Salinibacter* in this sample (Supplemental Figure S7), were similar to those annotated to Nocardioidaceae in the sediment sample, for example, heme biosynthesis, indicating a certain amount of overlap in non-core metabolic potential between the sediment and water. Others, like serotonin degradation and aromatic biogenic amine degradation, were unique to *Salinibacter*.

When comparing the abundance of 69 identified *Salinibacter*-associated pathways in the sediment versus the water (Supplemental Figure S7), a subset was identified in both samples (though at higher abundance in the water) and another portion that was almost entirely found in *Salinibacter* in the water. Interestingly, these aquatic *Salinibacter* specific pathways overlapped in function with many found in the sediment, most notably in the case of folate transformations, flavin biosynthesis, and chorismate biosynthesis.

### Lake Hillier shows BGC Potential

To determine the natural product potential of the Lake Hillier microbiome, analysis of Biosynthetic Gene Clusters (BGCs) was performed. AntiSMASH yielded a total of 129 BGCs across the two samples (Figure 6) for a broad range of small molecule classes, including polyketides (PKs), Terpene, Bacteriocin, Arylpolyene, Siderophore, Resorcinol, Nonribosomal peptide synthetase (NRPS)-like, among others. 127 of the 129 (98.4%) were novel, having less than 80% similarity with any categorized MIBiG entry of known secondary metabolites. Only 2 had an MIBiG reference (Terpene (BGC0000647) and Polyketide (BGC0001164)). Based on AntiSMASH, 3 of 51 distinct terpenes, including the one matching to BGC0000647, encoded a carotenoid biosynthetic gene. In total, from the current 1,926 known BGCs in the MiBiG database, less than 2% of the secondary metabolites found in our sediment and water samples of Lake Hillier are matched, giving potential to a novel source of new secondary metabolites.

**Figure 6:**
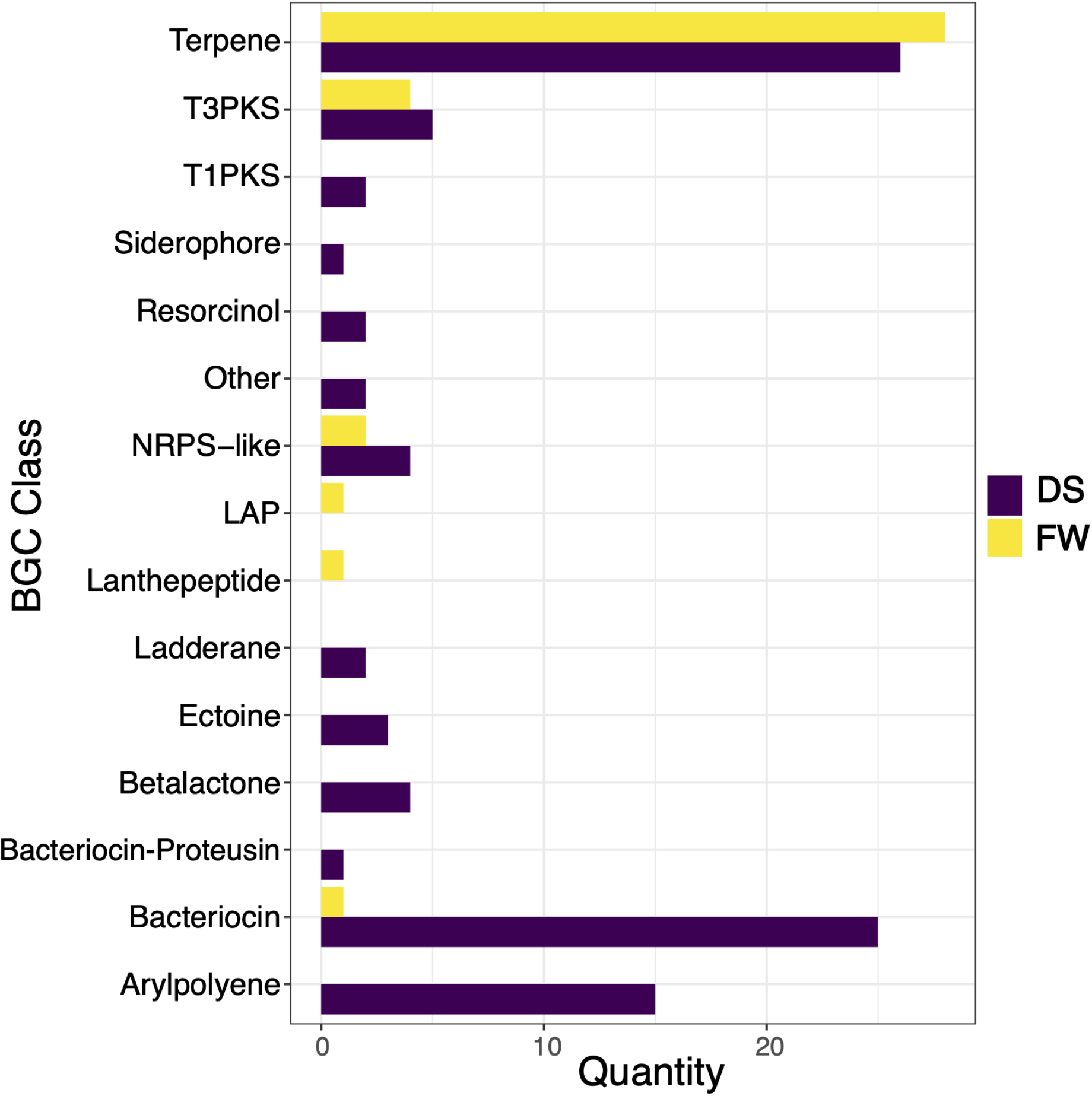
Biosynthetic Gene Clusters (BGCs) classes reported by AntiSMASH5 and their quantity in each one of the two metagenomes from sediment (DS) and water(FW).

Akin to the pathway-level analysis, the water and sediment samples encoded distinct BGC repertoires. Approximately 70% of total BGCs and also most of the BGC diversity was observed only in sediment samples. Two classes, Lanthepeptide and Linear azol(in)e-containing peptides (LAP) were only found in water. The most abundant BGCs classes identified were Terpene (both samples), Bacteriocin (mostly in sediment) and Arylpolyene (only in sediment). From the total observed 129 BGCs, there were 125 distinct families and of the 4 that clustered together, 3 were Terpenes and 1 encoded Ribosomally synthesized and post-translationally modified peptides (RiPPs). The remaining 121 samples did not cluster with each other.

### Culturing identifies organisms not highly abundant in sequencing-based approaches

Microscopic analysis of water samples revealed a high abundance of salt crystals, Figure S8A. A total of 13 isolates were recovered from sediments and water in both media. Culturing revealed a higher recovery of pigmented organisms on the Marine Broth Agar 2216 supplemented with NaCl (Figure S8B-D). Alignment and phylogenetic analysis showed that most isolates (n=5) belong to the genus *Bacillus*, Figure S8E, with support values ranging between 58% to 100%. However, none of the species from *Bacillus* that clustered with these isolates were found through WGS and amplicon sequence analysis. Similarly, the species *Aquibacillus halophilus* that clustered with one isolate (bootstrap 100%) was not identified by sequencing.

Other isolates grouped with members from genera *Jeotgalibacillus*, *Halobacillus*, and *Virgibacillus*, also found by WGS and amplicon sequencing. Lastly, one red-pigmented organism was recovered as the most abundant by culturing from water samples and assigned to *Psychroflexus tropicus* with 100% support, Figure S8E.

## Discussion

The biology of the deep pink color of Australia’s Lake Hillier, and of other ecosystems with similar coloration and phenotypes, has been a long-standing question. Here, we characterized the microbiome of this extreme ecosystem in an attempt to understand the potential sources for this unique color, and to determine any biochemically unique functions encoded by the organisms therein. We took a pan-domain, sequencing-based approach to characterize the archaea, bacteria, viruses, and algae present in Lake Hillier, identifying a preponderance of purple sulfur-producing microbes and a single metabolically dominant genus in the water: *Salinibacter*. The former microbes are associated with purple pigments, and the latter has been cultured and is known to be red-orange (66). As a result, we hypothesize that the color of Lake Hillier derives from a combination of these, and potentially other (e.g. *Dunaliella salina*), pigment-rich organisms.

These data demonstrate that Lake Hillier is composed of a diverse set of microorganisms ranging in four domains. In accordance with previous studies (44; 87), these results evidence the poor correlation between sequencing methods (shotgun vs. amplicon), where only a few subsets of species overlapped between the two (Figure 2). Comparison of preservation methods did not reveal a significant difference in abundances which has been a long pondered question among extremophiles researchers (88). This may suggest that for an ecosystem of extreme salt concentration microbial durability may be more consistent and unaffected by chemical exposures. Furthermore, these analyses described a consistent set of 53 microorganisms among all samples that denominate the Lake Hillier *core* microbiome, but only a subset of 37 taxa was classified to species level. Core species have been reported in different hosts (2; 36; 79) and environments (20; 22; 68) as functional contributors to the host survival and steady state of the environment. Most of Lake Hillier core species have been previously isolated from saline lakes in Senegal, Russia, Egypt, Spain (4; 63; 81; 83), solar salterns (19; 91; 96), contaminated environments (48; 50), and as algae symbionts (92). However, the taxonomic identity of half of the core species in Lake Hillier remains to be assigned.

These metagenomic analyses also detected 112 viral taxa, many associated with Haloviruses which have been previously reported in hypersaline environments (5; 80) and in the Yellowstone Lake (101). Consistent with previous studies of hypersaline environments, archaea viruses of order Caudovirales had the most representatives in these data. (51; 76). A few taxa of “giant viruses” from orders Algavirales, Chitovirales, Pimascovirales and Imitervirales, were also identified, which are known to infect a wide range of algae in marine environments (6; 33). Although these findings are far from representing the true diversity of Lake Hillier virome, as it is estimated that millions of viruses can be found in one milliliter of ocean water (85; 97), these results corroborate previous studies from hypersaline environments and highlight the need for additional metagenomic studies to describe and understand the viral dynamics and influence on the microbiome.

Microbial profiling exhibited the presence of almost 500 extremophiles in the lake, and this number is likely to be underestimated due to the ongoing expansion of the Microbe Directory (TMD) (78). Though useful, TMD is almost certainly an incomplete list of microbes and characteristics, as it is based on manual curation of published articles. Nevertheless, this profiling serves as a first glimpse of the diversity of polyextremophiles in Lake Hillier and any other extreme environments. Additionally, a considerable diversity of purple-bacteria, with sulfur and non-sulfur capabilities, were identified and subsequently consolidate in our pathways analysis.

Functionally redundant, yet taxonomically discrete pathways were identified between the sediment and water WGS samples. While the highly abundant fundamental metabolisms between these two micro-environments were similar, they are housed in two phylogenetically distinct and dominant groups: *Salinibacter ruber* in the water, and an unclassified *Nocardiodaceae bacterium* in the sediment. The latter is known to be wide ranging in terms of ecology (e.g. from lakes to soil to the built environment) and potentially carry a number of functions for bioremediation of extreme or polluted habitats (7; 31; 75). Notably, we were able to extract partial MAGs for representative strains of both *Salinibacter ruber* and *Nocardiodaceae* organisms. Overall, microbes from Lake Hillier displayed a variable metabolic capacity, where mostly universal pathways were similarly prevalent in sediments and water, while more specialized pathways depended on the sample type.

More relevant to natural product discovery, these data clearly demonstrate the capacity of Lake Hillier’s microbiome to produce a variety of uncommon and uncharacterized secondary metabolites. Microbial secondary metabolites are of great biotechnological interest (29). Only until recently, the ecological relevance of small molecules within the microbiomes has been displayed, where complex biological systems exhibit equally complex metabolites arsenal (23; 55). While there are only a few studies looking at the prevalence of BGCs in extreme environments (11; 18; 72), new machine learning approaches have evidenced the influence of extreme conditions (e.i. saline concentrations) in the production of secondary metabolites in halophiles (43). Although recent computational approaches provide an advantage to identify active and silent BGCs (90), further analyses are needed to characterize which of the 129 BGCs identified in Lake Hillier actively encode secondary metabolites.

Hypersaline environments have previously been a source of novel metagenome-assembled genomes (MAGs) (7; 100). The binning approaches described here resulted in partial, and in some cases, nearly complete genomes that spanned the tree of life (Figure 4C) and were taxonomically annotated to similar clades as many organisms in the short read and amplicon analysis. Instead of applying hard cutoffs to eliminate low quality but interesting genomes, we opted to include them in the analysis as they may serve as potential indicators of where we could achieve complete genomes in future studies. One potential drawback of this approach is that the lack of ability to annotate bins to the species level could be ascribed to a high proportion of novel genomes in Lake Hillier, low quality or high contamination in genome bins, or a combination of the two. Regardless, based on these analyses, specifically our ability to recover at least partial genomes for abundant and biochemically unique organisms (e.g. *Salinibacter*) we hypothesize further sequencing depth and additional methods (e.g. long-reads) would yield increased novel genomes with valuable biomining potential.

While sequencing and culturing of Lake Hillier samples do not completely concur, previous studies have attributed this discrepancy to the microbes specificity to certain culture media, temperatures, pH, growth rates, among other factors (38). In metagenomic analysis, the high salt concentrations of Lake Hillier, insufficient lysis, and poor DNA recovery may have contributed to this difference from culture based methods, which has been previously evidenced in other extreme environments (37). Further characterization of the metabolic requirements of microbes from Lake Hillier and other extreme microbiomes is needed to design specific culture-based approaches to target the broad diversity of this hypersaline environment.

## Conclusions

Overall, we do not claim here to have the final answer on the source of Lake Hillier’s color – indeed, a deeper, likely culture-based, exploration of the organisms and metabolisms identified here will likely be necessary to definitively address this question. However, we propose the exploration of *Salinibacter* and the other pigment-producing organisms identified here as the first step in these future experiments. We additionally expect that further sequencing, culturing, and comprehensive microscopy of Lake Hillier and other similar ecosystems will provide a wellspring of biotechnological potential and extremophilic biology that can only be found in these idiosyncratic environments.

## Supporting information

Sierra et al Supplemental Data

## Acknowledgements

The authors would like to thank the support of the Western Australia Department of Parks and Wildlife (DPaW), and John Lizamore and Don Cater for permits and assistance with sampling the lake, as well as the Australian Genome Research Facility (AGRF) and Bioplatforms Australia for infrastructure support. We would like to thank Phil Hugenholtz (University of Queensland) for supporting the expedition to Lake Hillier. Additionally, RNA sequencing was provided by Weill Cornell Medicine, with the support of Jorge Gandara, which was ultimately excluded from the study. We would like to recognize Audria Greenwald (UVM) for their processing and DNA extractions of the water and sediment samples from Lake Hillier. We also acknowledge Jaden J. A. Hastings (WCM) and Deena Najjar (WCM) for their valued input on the project. Additionally, Illumina, Inc. is acknowledged for providing sequencing support for the WGS and RNA-seq data.

## Ethical Approval and Consent to participate

“Not applicable”

## Abbreviations

XMP: Extreme Microbiome Project
AGRF: Australian Genome Research Facility
WGS: Whole Genome Sequencing
TMD: The Microbe Directory
GTDB: Genome Taxonomy Database
PSB: Purple sulfur-bacteria
NPSN: Purple non-sulfur-bacteria
MAGs: Metagenome-Assembled Genomes
ANI: Average Nucleotide Identity
BGCs: Biosynthetic Gene Clusters
GCF: Gene Cluster Family

## Availability of data and materials

Sequences and scripts will be made available upon publication.

## Consent for publication

All authors read and approved this manuscript.

## Authors’ contributions

The project originated under S.W.T., C.E.M., and K.M. with assistance from M.G.R.G. K.M, and S.W.T obtained funding, logistics, and permits for sampling. K.M. conducted all field sampling. S.W.T. and K.M performed DNA extractions. S.W.T., K.M, S.G., W.K.T., and J.R. performed library preparation, sequencing, and initial analyses. N.J.B conducted full length 16S rRNA sequencing on pure cultures. S.W.T performed microscopy and culturing. D.B. coordinated data storage. K.A.R. provided input for methods and materials. M.A.S.,B.T.T.,J.F., E.A., and C.B. analyzed the data. M.A.S., B.T.T. S.W.T, K.M, and K.A.R wrote the manuscript. All authors read and approved the final manuscript.

## Authors’ information

Maria A. Sierra, Krista A. Ryon and Braden T. Tierney contributed equally to this work.

## References

[1] (2015). Fastqc.

[2] Ainsworth, T. D., Krause, L., Bridge, T., Torda, G., Raina, J.-B., Zakrzewski, M., Gates, R. D., Padilla-Gamiño, J. L., Spalding, H. L., Smith, C., et al. (2015). The coral core microbiome identifies rare bacterial taxa as ubiquitous endosymbionts. The ISME journal, 9(10):2261.

[3] Alneberg, J., Bjarnason, B. S., de Bruijn, I., Schirmer, M., Quick, J., Ijaz, U. Z., Loman, N. J., Andersson, A. F., and Quince, C. (2013). Concoct: clustering contigs on coverage and composition. arXiv preprint arXiv:1312.4038.

[4] Antón, J., Oren, A., Benlloch, S., Rodríguez-Valera, F., Amann, R., and Rosselló-Mora, R. (2002). Salinibacter ruber gen. nov., sp. nov., a novel, extremely halophilic member of the bacteria from saltern crystallizer ponds. International journal of systematic and evolutionary microbiology, 52(2):485–491.

[5] Atanasova, N. S., Oksanen, H. M., and Bamford, D. H. (2015). Haloviruses of archaea, bacteria, and eukaryotes. Current opinion in microbiology, 25:40–48.

[6] Aylward, F., Moniruzzaman, M., Ha, A. D., and Koonin, E. V. (2021). A phylogenomic framework for charting the diversity and evolution of giant viruses. bioRxiv.

[7] Bardavid, R. E. and Oren, A. (2008). Dihydroxyacetone metabolism in salinibacter ruber and in haloquadratum walsbyi. Extremophiles, 12(1):125–131.

[8] Becker, E. A., Seitzer, P. M., Tritt, A., Larsen, D., Krusor, M., Yao, A. I., Wu, D., Madern, D., Eisen, J. A., Darling, A. E., et al. (2014). Phylogenetically driven sequencing of extremely halophilic archaea reveals strategies for static and dynamic osmo-response. PLoS genetics, 10(11):e1004784.

[9] Beghini, F., McIver, L. J., Blanco-Míguez, A., Dubois, L., Asnicar, F., Maharjan, S., Mailyan, A., Manghi, P., Scholz, M., Thomas, A. M., et al. (2021). Integrating taxonomic, functional, and strain-level profiling of diverse microbial communities with biobakery 3. Elife, 10:e65088.

[10] Bermanec, V., Paradžik, T., Kazazić, S. P., Venter, C., Hrenović, J., Vujaklija, D., Duran, R., Boev, I., and Boev, B. (2021). Novel arsenic hyper-resistant bacteria from an extreme environment, crven dol mine, allchar, north macedonia. Journal of Hazardous Materials, 402:123437.

[11] Bhattacharya, C. (2020). Decoding the Cryptic Metagenome: A Deep Dive into Gene Clusters and Taxonomy of Microbiome. PhD thesis, Weill Medical College of Cornell University.

[12] Blin, K., Shaw, S., Steinke, K., Villebro, R., Ziemert, N., Lee, S. Y., Medema, M. H., and Weber, T. (2019). antismash 5.0: updates to the secondary metabolite genome mining pipeline. Nucleic acids research, 47(W1):W81–W87.

[13] Bolyen, E., Rideout, J. R., Dillon, M. R., Bokulich, N. A., Abnet, C. C., Al-Ghalith, G. A., Alexander, H., Alm, E. J., Arumugam, M., Asnicar, F., et al. (2019). Reproducible, interactive, scalable and extensible microbiome data science using qiime 2. Nature biotechnology, 37(8):852–857.

[14] Brock, T. D. and Freeze, H. (1969). Thermus aquaticus gen. n. and sp. n., a nonsporulating extreme thermophile. Journal of bacteriology, 98(1):289–297.

[15] Bushnell, B. (2014). Bbmap: a fast, accurate, splice-aware aligner. Technical report, Lawrence Berkeley National Lab.(LBNL), Berkeley, CA (United States).

[16] Çelebi, H., Bahadır, T., Şimşek, İ., and Tulun, Ş. (2021). Use of dunaliella salina in environmental applications.

[17] Chaumeil, P.-A., Mussig, A. J., Hugenholtz, P., and Parks, D. H. (2020). Gtdb-tk: a toolkit to classify genomes with the genome taxonomy database.

[18] Chen, R., Wong, H. L., Kindler, G. S., MacLeod, F. I., Benaud, N., Ferrari, B. C., and Burns, B. P. (2020). Discovery of an abundance of biosynthetic gene clusters in shark bay microbial mats. Frontiers in Microbiology, 11:1950.

[19] Chen, S., Xu, Y., Liu, H.-C., Yang, A.-N., and Ke, L.-X. (2017). Halobaculum roseum sp. nov., isolated from underground salt deposits. International journal of systematic and evolutionary microbiology, 67(4):818–823.

[20] Danko, D., Bezdan, D., Afshin, E. E., Ahsanuddin, S., Bhattacharya, C., Butler, D. J., Chng, K. R., Donnellan, D., Hecht, J., Jackson, K., et al. (2021). A global metagenomic map of urban microbiomes and antimicrobial resistance. Cell.

[21] DasSarma, S. and DasSarma, P. (2015). Halophiles and their enzymes: negativity put to good use. Current opinion in microbiology, 25:120–126.

[22] Di Gregorio, L., Tandoi, V., Congestri, R., Rossetti, S., and Di Pippo, F. (2017). Unravelling the core microbiome of biofilms in cooling tower systems. Biofouling, 33(10):793–806.

[23] Donia, M. S., Cimermancic, P., Schulze, C. J., Brown, L. C. W., Martin, J., Mitreva, M., Clardy, J., Linington, R. G., and Fischbach, M. A. (2014). A systematic analysis of biosynthetic gene clusters in the human microbiome reveals a common family of antibiotics. Cell, 158(6):1402–1414.

[24] Edgar, R. C. (2004). Muscle: multiple sequence alignment with high accuracy and high throughput. Nucleic acids research, 32(5):1792–1797.

[25] Edwards, K. J., Bond, P. L., Gihring, T. M., and Banfield, J. F. (2000). An archaeal iron-oxidizing extreme acidophile important in acid mine drainage. Science, 287(5459):1796–1799.

[26] Emerson, J. B., Andrade, K., Thomas, B. C., Norman, A., Allen, E. E., Heidelberg, K. B., and Banfield, J. F. (2013a). Virus-host and crispr dynamics in archaea-dominated hypersaline lake tyrrell, victoria, australia. Archaea, 2013.

[27] Emerson, J. B., Thomas, B. C., Andrade, K., Heidelberg, K. B., and Banfield, J. F. (2013b). New approaches indicate constant viral diversity despite shifts in assemblage structure in an australian hypersaline lake. Applied and environmental microbiology, 79(21):6755–6764.

[28] Empadinhas, N. and da Costa, M. S. (2011). Diversity, biological roles and biosynthetic pathways for sugar-glycerate containing compatible solutes in bacteria and archaea. Environmental microbiology, 13(8):2056–2077.

[29] Giordano, D., Coppola, D., Russo, R., Denaro, R., Giuliano, L., Lauro, F. M., di Prisco, G., and Verde, C. (2015). Marine microbial secondary metabolites: pathways, evolution and physiological roles. Advances in microbial physiology, 66:357–428.

[30] Gloor, G. (2015). Aldex2: Anova-like differential expression tool for compositional data. ALDEX manual modular, 20:1–11.

[31] Goodfellow, M. (2014). The family nocardiaceae. The Prokaryotes: Actinobacteria.

[32] Gray, D. A., Dugar, G., Gamba, P., Strahl, H., Jonker, M. J., and Hamoen, L. W. (2019). Extreme slow growth as alternative strategy to survive deep starvation in bacteria. Nature communications, 10(1):1–12.

[33] Ha, A. D., Moniruzzaman, M., and Aylward, F. O. (2021). High transcriptional activity and diverse functional repertoires of hundreds of giant viruses in a coastal marine system. bioRxiv.

[34] Harding, T., Roger, A. J., and Simpson, A. G. (2017). Adaptations to high salt in a halophilic protist: differential expression and gene acquisitions through duplications and gene transfers. Frontiers in microbiology, 8:944.

[35] Heidelberg, K. B., Nelson, W. C., Holm, J. B., Eisenkolb, N., Andrade, K., and Emerson, J. B. (2013). Characterization of eukaryotic microbial diversity in hypersaline lake tyrrell, australia. Frontiers in Microbiology, 4:115.

[36] Henderson, G., Cox, F., Ganesh, S., Jonker, A., Young, W., and Janssen, P. H. (2015). Rumen microbial community composition varies with diet and host, but a core microbiome is found across a wide geographical range. Scientific reports, 5(1):1–15.

[37] Herrera, A. and Cockell, C. S. (2007). Exploring microbial diversity in volcanic environments: a review of methods in dna extraction. Journal of Microbiological Methods, 70(1):1–12.

[38] Hiergeist, A., Gläsner, J., Reischl, U., and Gessner, A. (2015). Analyses of intestinal microbiota: culture versus sequencing. ILAR journal, 56(2):228–240.

[39] Horita, J. (2005). Saline waters. In Isotopes in the water cycle, pages 271–287. Springer.

[40] Huerta-Cepas, J., Serra, F., and Bork, P. (2016). Ete 3: reconstruction, analysis, and visualization of phylogenomic data. Molecular biology and evolution, 33(6):1635–1638.

[41] Hurst, C. J. (2016). Their World: A diversity of microbial environments, volume 1. Springer.

[42] Hyatt, D., Chen, G.-L., LoCascio, P. F., Land, M. L., Larimer, F. W., and Hauser, L. J. (2010). Prodigal: prokaryotic gene recognition and translation initiation site identification. BMC bioinformatics, 11(1):1–11.

[43] Jančič, S., Frisvad, J. C., Kocev, D., Gostinčar, C., Džeroski, S., and Gunde-Cimerman, N. (2016). Production of secondary metabolites in extreme environments: Food-and airborne wallemia spp. produce toxic metabolites at hypersaline conditions. PLoS One, 11(12):e0169116.

[44] Jovel, J., Patterson, J., Wang, W., Hotte, N., O’Keefe, S., Mitchel, T., Perry, T., Kao, D., Mason, A. L., Madsen, K. L., et al. (2016). Characterization of the gut microbiome using 16s or shotgun metagenomics. Frontiers in microbiology, 7:459.

[45] Kang, D. D., Froula, J., Egan, R., and Wang, Z. (2015). Metabat, an efficient tool for accurately reconstructing single genomes from complex microbial communities. PeerJ, 3:e1165.

[46] Kautsar, S. A., Blin, K., Shaw, S., Navarro-Muñoz, J. C., Terlouw, B. R., van der Hooft, J. J., Van Santen, J. A., Tracanna, V., Suarez Duran, H. G., Pascal Andreu, V., et al. (2020). Mibig 2.0: a repository for biosynthetic gene clusters of known function. Nucleic acids research, 48(D1):D454–D458.

[47] Kautsar, S. A., Blin, K., Shaw, S., Weber, T., and Medema, M. H. (2021). Big-fam: the biosynthetic gene cluster families database. Nucleic acids research, 49(D1):D490–D497.

[48] Kleindienst, S., Herbst, F.-A., Stagars, M., Von Netzer, F., Von Bergen, M., Seifert, J., Peplies, J., Amann, R., Musat, F., Lueders, T., et al. (2014). Diverse sulfate-reducing bacteria of the desulfosarcina/desulfococcus clade are the key alkane degraders at marine seeps. The ISME journal, 8(10):2029–2044.

[49] Larsson, A. (2014). Aliview: a fast and lightweight alignment viewer and editor for large datasets. Bioinformatics, 30(22):3276–3278.

[50] Li, P., Li, B., Webster, G., Wang, Y., Jiang, D., Dai, X., Jiang, Z., Dong, H., and Wang, Y. (2014). Abundance and diversity of sulfate-reducing bacteria in high arsenic shallow aquifers. Geomicrobiology Journal, 31(9):802–812.

[51] Liu, Y., Demina, T. A., Roux, S., Aiewsakun, P., Kazlauskas, D., Simmonds, P., Prangishvili, D., Oksanen, H. M., and Krupovic, M. (2021). Diversity, taxonomy and evolution of archaeal viruses of the class caudoviricetes. bioRxiv.

[Lizamore] Lizamore, J. Water quality review of pink lake and associated lakes.

[53] Lu, J., Breitwieser, F. P., Thielen, P., and Salzberg, S. L. (2017). Bracken: estimating species abundance in metagenomics data. PeerJ Computer Science, 3:e104.

[54] Maria Sierra David Danko (2020). The microbe directory.

[55] Medina, A., Schmidt-Heydt, M., Rodriguez, A., Parra, R., Geisen, R., and Magan, N. (2015). Impacts of environmental stress on growth, secondary metabolite biosynthetic gene clusters and metabolite production of xerotolerant/xerophilic fungi. Current genetics, 61(3):325–334.

[56] Merroun, M. L. and Selenska-Pobell, S. (2008). Bacterial interactions with uranium: an environmental perspective. Journal of Contaminant Hydrology, 102(3-4):285–295.

[57] Mikheenko, A., Saveliev, V., and Gurevich, A. (2016). Metaquast: evaluation of metagenome assemblies. Bioinformatics, 32(7):1088–1090.

[58] Mormile, M. R., Hong, B.-Y., and Benison, K. C. (2009). Molecular analysis of the microbial communities of mars analog lakes in western australia. Astrobiology, 9(10):919–930.

[59] Navarro-Muñoz, J. C., Selem-Mojica, N., Mullowney, M. W., Kautsar, S., Tryon, J. H., Parkinson, E. I., De Los Santos, E. L., Yeong, M., Cruz-Morales, P., Abubucker, S., et al. (2018). A computational framework for systematic exploration of biosynthetic diversity from large-scale genomic data. Biorxiv, page 445270.

[60] Nguyen, L.-T., Schmidt, H. A., Von Haeseler, A., and Minh, B. Q. (2015). Iq-tree: a fast and effective stochastic algorithm for estimating maximum-likelihood phylogenies. Molecular biology and evolution, 32(1):268–274.

[61] Norambuena, J. (2020). Mechanism of resistance focusing on copper, mercury and arsenic in extremophilic organisms, how acidophiles and thermophiles cope with these metals. In Physiological and Biotechnological Aspects of Extremophiles, pages 23–37. Elsevier.

[62] Nurk, S., Meleshko, D., Korobeynikov, A., and Pevzner, P. A. (2017). metaspades: a new versatile metagenomic assembler. Genome research, 27(5):824–834.

[63] Ollivier, B., Hatchikian, C., Prensier, G., Guezennec, J., and Garcia, J.-L. (1991). Desulfohalobium retbaense gen. nov., sp. nov., a halophilic sulfate-reducing bacterium from sediments of a hypersaline lake in senegal. International Journal of Systematic and Evolutionary Microbiology, 41(1):74–81.

[64] Olm, M. R., Brown, C. T., Brooks, B., and Banfield, J. F. (2017). drep: a tool for fast and accurate genomic comparisons that enables improved genome recovery from metagenomes through de-replication. The ISME journal, 11(12):2864–2868.

[65] Ondov, B. D., Starrett, G. J., Sappington, A., Kostic, A., Koren, S., Buck, C. B., and Phillippy, A. M. (2019). Mash screen: high-throughput sequence containment estimation for genome discovery. Genome biology, 20(1):1–13.

[66] Oren, A. (2015). Salinibacter: an extremely halophilic bacterium with archaeal properties. FEMS Microbiology Letters, 342(1):1–9.

[67] Parks, D. H., Imelfort, M., Skennerton, C. T., Hugenholtz, P., and Tyson, G. W. (2015). Checkm: assessing the quality of microbial genomes recovered from isolates, single cells, and metagenomes. Genome research, 25(7):1043–1055.

[68] Pershina, E. V., Ivanova, E. A., Korvigo, I. O., Chirak, E. L., Sergaliev, N. H., Abakumov, E. V., Provorov, N. A., and Andronov, E. E. (2018). Investigation of the core microbiome in main soil types from the east european plain. Science of the Total Environment, 631:1421–1430.

[69] Podell, S., Emerson, J. B., Jones, C. M., Ugalde, J. A., Welch, S., Heidelberg, K. B., Banfield, J. F., and Allen, E. E. (2014). Seasonal fluctuations in ionic concentrations drive microbial succession in a hypersaline lake community. The ISME journal, 8(5):979–990.

[70] Porter, K., Kukkaro, P., Bamford, J. K., Bath, C., Kivelä, H. M., Dyall-Smith, M. L., and Bamford, D. H. (2005). Sh1: a novel, spherical halovirus isolated from an australian hypersaline lake. Virology, 335(1):22–33.

[71] Raddadi, N., Cherif, A., Daffonchio, D., Neifar, M., and Fava, F. (2015). Biotechnological applications of extremophiles, extremozymes and extremolytes. Applied microbiology and biotechnology, 99(19):7907–7913.

[72] Rego, A., Fernandez-Guerra, A., Duarte, P., Assmy, P., Leão, P. N., and Magalhães, C. (2021). Secondary metabolite biosynthetic diversity in arctic ocean metagenomes. Microbial Genomics, 7(12):000731.

[73] Reysenbach, A. (1995). Archaea: A laboratory manual thermophiles. CSHLP, pages 101–107.

[74] Robeson, M. S., O’Rourke, D. R., Kaehler, B. D., Ziemski, M., Dillon, M. R., Foster, J. T., and Bokulich, N. A. (2021). Rescript: Reproducible sequence taxonomy reference database management. PLoS computational biology, 17(11):e1009581.

[75] Rossello-Mora, R., Lucio, M., Pena, A., Brito-Echeverría, J., Lopez-Lopez, A., Valens-Vadell, M., Frommberger, M., Anton, J., and Schmitt-Kopplin, P. (2008). Metabolic evidence for biogeographic isolation of the extremophilic bacterium salinibacter ruber. The ISME journal, 2(3):242–253.

[76] Roux, S., Enault, F., Ravet, V., Colombet, J., Bettarel, Y., Auguet, J.-C., Bouvier, T., Lucas-Staat, S., Vellet, A., Prangishvili, D., et al. (2016). Analysis of metagenomic data reveals common features of halophilic viral communities across continents. Environmental microbiology, 18(3):889–903.

[77] Schönknecht, G., Chen, W.-H., Ternes, C. M., Barbier, G. G., Shrestha, R. P., Stanke, M., Bräutigam, A., Baker, B. J., Banfield, J. F., Garavito, R. M., et al. (2013). Gene transfer from bacteria and archaea facilitated evolution of an extremophilic eukaryote. Science, 339(6124):1207–1210.

[78] Sierra, M. A., Bhattacharya, C., Ryon, K., Meierovich, S., Shaaban, H., Westfall, D., Mohammad, R., Kuchin, K., Afshinnekoo, E., Danko, D. C., et al. (2019). The microbe directory v2. 0: An expanded database of ecological and phenotypical features of microbes. BioRxiv.

[79] Sierra, M. A., Danko, D. C., Sandoval, T. A., Pishchany, G., Moncada, B., Kolter, R., Mason, C. E., and Zambrano, M. M. (2020). The microbiomes of seven lichen genera reveal host specificity, a reduced core community and potential as source of antimicrobials. Frontiers in microbiology, 11:398.

[80] Sime-Ngando, T., Lucas, S., Robin, A., Tucker, K. P., Colombet, J., Bettarel, Y., Desmond, E., Gribaldo, S., Forterre, P., Breitbart, M., et al. (2011). Diversity of virus–host systems in hypersaline lake retba, senegal. Environmental Microbiology, 13(8):1956–1972.

[81] Singh, K. S., Kirksey, J., Hoff, W. D., and Deole, R. (2014). Draft genome sequence of the extremely halophilic phototrophic purple sulfur bacterium halorhodospira halochloris. Journal of genomics, 2:118.

[82] Sogin, M. L. (1990). Amplification of rrna genes for molecular evolution studies. PCR Protocols: A Guide to Methods and Applications, pages 307–314.

[83] Sorokin, D. Y., Messina, E., Smedile, F., Roman, P., Damsté, J. S. S., Ciordia, S., Mena, M. C., Ferrer, M., Golyshin, P. N., Kublanov, I. V., et al. (2017). Discovery of anaerobic lithoheterotrophic haloarchaea, ubiquitous in hypersaline habitats. The ISME journal, 11(5):1245–1260.

[84] Stan-Lotter, H. and Fendrihan, S. (2012). Adaption of microbial life to environmental extremes. Springer.

[85] Suttle, C. A. (2007). Marine viruses—major players in the global ecosystem. Nature reviews microbiology, 5(10):801–812.

[86] Takayanagi, S., Kawasaki, H., Sugimori, K., Yamada, T., Sugai, A., Ito, T., Yamasato, K., and Shioda, M. (1996). Sulfolobus hakonensis sp. nov., a novel species of acidothermophilic archaeon. International Journal of Systematic and Evolutionary Microbiology, 46(2):377–382.

[87] Tessler, M., Neumann, J. S., Afshinnekoo, E., Pineda, M., Hersch, R., Velho, L. F. M., Segovia, B. T., Lansac-Toha, F. A., Lemke, M., DeSalle, R., et al. (2017). Large-scale differences in microbial biodiversity discovery between 16s amplicon and shotgun sequencing. Scientific reports, 7(1):1–14.

[88] Tighe, S., Afshinnekoo, E., Rock, T. M., McGrath, K., Alexander, N., McIntyre, A., Ahsanuddin, S., Bezdan, D., Green, S. J., Joye, S., et al. (2017). Genomic methods and microbiological technologies for profiling novel and extreme environments for the extreme microbiome project (xmp). Journal of biomolecular techniques: JBT, 28(1):31.

[89] Uritskiy, G. V., DiRuggiero, J., and Taylor, J. (2018). Metawrap—a flexible pipeline for genome-resolved metagenomic data analysis. Microbiome, 6(1):1–13.

[90] van der Lee, T. A. and Medema, M. H. (2016). Computational strategies for genome-based natural product discovery and engineering in fungi. Fungal Genetics and Biology, 89:29–36.

[91] Vreeland, R. H., Litchfield, C., Martin, E., and Elliot, E. (1980). Halomonas elongata, a new genus and species of extremely salt-tolerant bacteria. International Journal of Systematic and Evolutionary Microbiology, 30(2):485–495.

[92] Wang, G., Tang, M., Li, T., Dai, S., Wu, H., Chen, C., He, H., Fan, J., Xiang, W., and Li, X. (2015). Wenzhouxiangella marina gen. nov, sp. nov, a marine bacterium from the culture broth of picochlorum sp. 122, and proposal of wenzhouxiangellaceae fam. nov. in the order chromatiales. Antonie Van Leeuwenhoek, 107(6):1625–1632.

[93] Wibowo, M. C., Yang, Z., Borry, M., Hübner, A., Huang, K. D., Tierney, B. T., Zimmerman, S., Barajas-Olmos, F., Contreras-Cubas, C., García-Ortiz, H., et al. (2021). Reconstruction of ancient microbial genomes from the human gut. Nature, 594(7862):234–239.

[94] Wood, D. E., Lu, J., and Langmead, B. (2019). Improved metagenomic analysis with kraken 2. Genome biology, 20(1):1–13.

[95] Wu, Y.-W., Simmons, B. A., and Singer, S. W. (2016). Maxbin 2.0: an automated binning algorithm to recover genomes from multiple metagenomic datasets. Bioinformatics, 32(4):605–607.

[96] Xia, J., Zhao, J.-X., Sang, J., Chen, G.-J., and Du, Z.-J. (2017). Halofilum ochraceum gen. nov., sp. nov., a gammaproteobacterium isolated from a marine solar saltern. International journal of systematic and evolutionary microbiology, 67(4):932–938.

[97] Yau, S., Lauro, F. M., DeMaere, M. Z., Brown, M. V., Thomas, T., Raftery, M. J., Andrews-Pfannkoch, C., Lewis, M., Hoffman, J. M., Gibson, J. A., et al. (2011). Virophage control of antarctic algal host–virus dynamics. Proceedings of the National Academy of Sciences, 108(15):6163–6168.

[98] Yu, G. (2020). Using ggtree to visualize data on tree-like structures. Current protocols in bioinformatics, 69(1):e96.

[99] Yu, G., Smith, D. K., Zhu, H., Guan, Y., and Lam, T. T.-Y. (2017). ggtree: an r package for visualization and annotation of phylogenetic trees with their covariates and other associated data. Methods in Ecology and Evolution, 8(1):28–36.

[100] Zhao, D., Zhang, S., Xue, Q., Chen, J., Zhou, J., Cheng, F., Li, M., Zhu, Y., Yu, H., Hu, S., et al. (2020). Abundant taxa and favorable pathways in the microbiome of soda-saline lakes in inner mongolia. Frontiers in microbiology, 11:1740.

[101] Zhou, J., Sun, D., Childers, A., McDermott, T. R., Wang, Y., and Liles, M. R. (2015). Three novel virophage genomes discovered from yellowstone lake metagenomes. Journal of virology, 89(2):1278–1285.

