## Supplementary material for "Cross-kingdom metagenomic profiling of Lake Hillier reveals pigment-rich polyextremophiles and wide-ranging metabolic adaptations": Sierra et al Supplemental Data

### List Appendix

#### Supplemental Table 1.

Summary statistics on assemblies and bins

#### Supplemental Figure 1.

Images of Lake Hillier from date of sample collection.

#### Supplemental Figure 2.

Relative abundance of taxa at the phylum level for Bacteria from different sample types. Size of dots represents abundance of taxon.

#### Supplemental Figure 3.

Relative abundance of taxa at the phylum level for Archaea (A), Eukaryota (B) and Viruses (C) from different sample types. Size of dots represents abundance of taxon.

#### Supplemental Figure 4.

Number of reads of top 20 most abundant species found by amplicon and whole genome sequencing (WGS) sequencing methods in Bacteria (A-B), Archaea(C-D), Eukaryotes(E,F), Virus(G).

#### Supplemental Figure 5.

Species overlap between sample types: Water, Bank and Sediment

#### Supplemental Figure 6.

Number of reads of Purple sulfur and non-sulfur bacteria present in Bank, Sediment and Water.

#### Supplemental Figure 7.

Difference in pathways from the two metagenomes from water (FW) and sediment (DS) in *Salinibater.*

#### Supplemental Figure 8.

Cultures of sediment and water samples from Lake Hillier.

#### Supplemental Figures

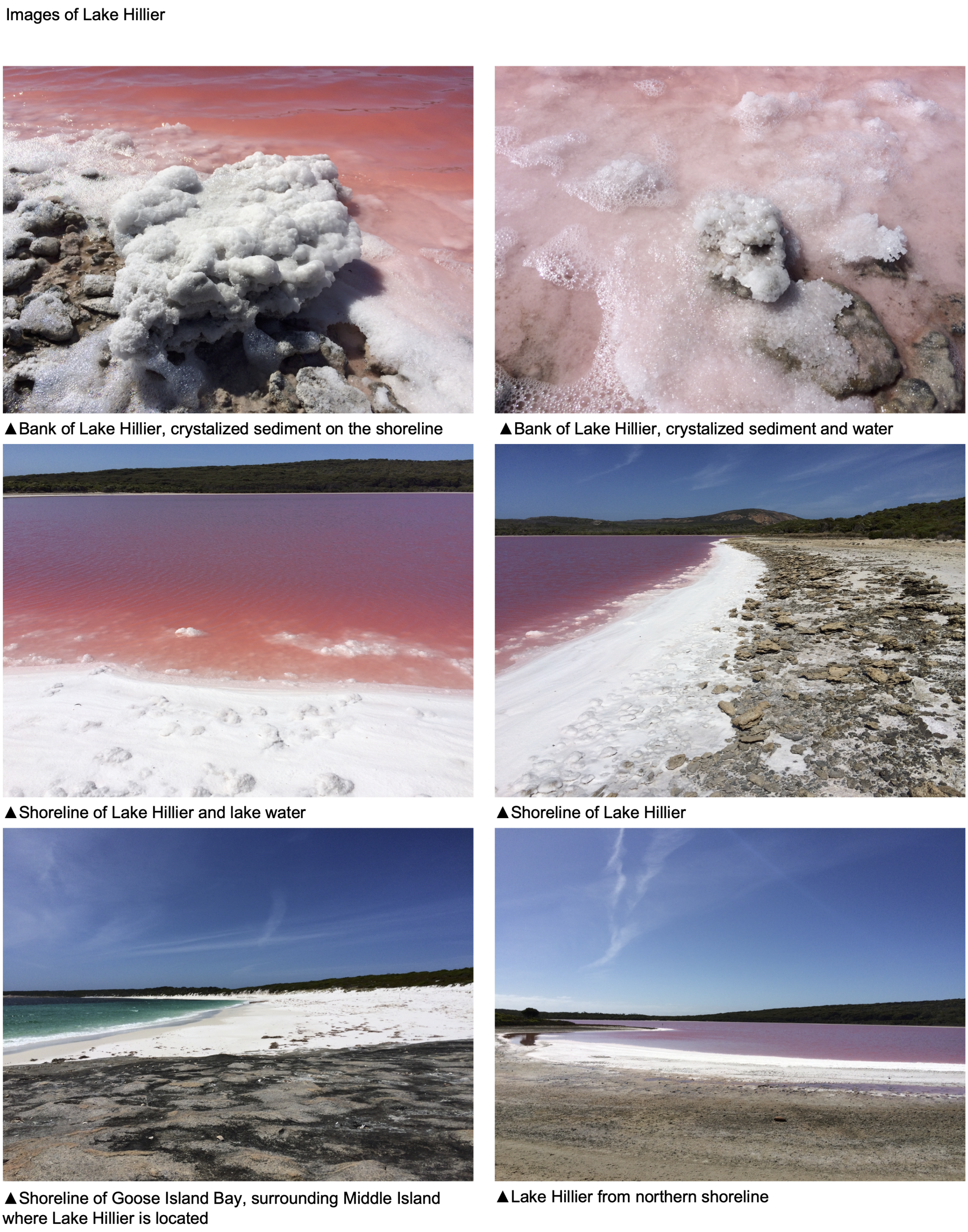


Figure S1: Pictures of Lake Hillier taken during the day of sample collection.

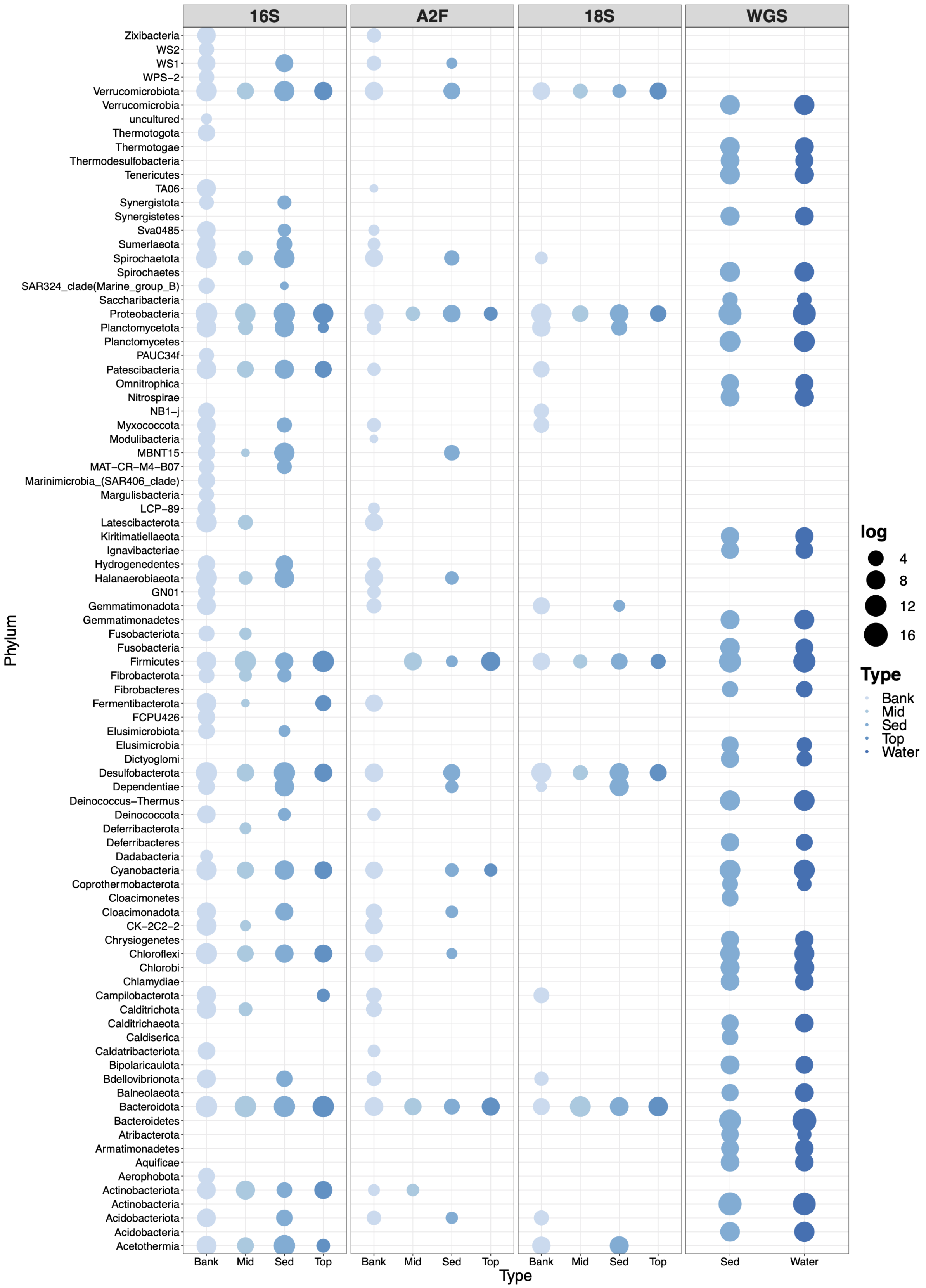


Figure S2: Relative abundance of taxa at the phylum level for Bacteria from different sample types. Size of dots represents abundance of taxon.

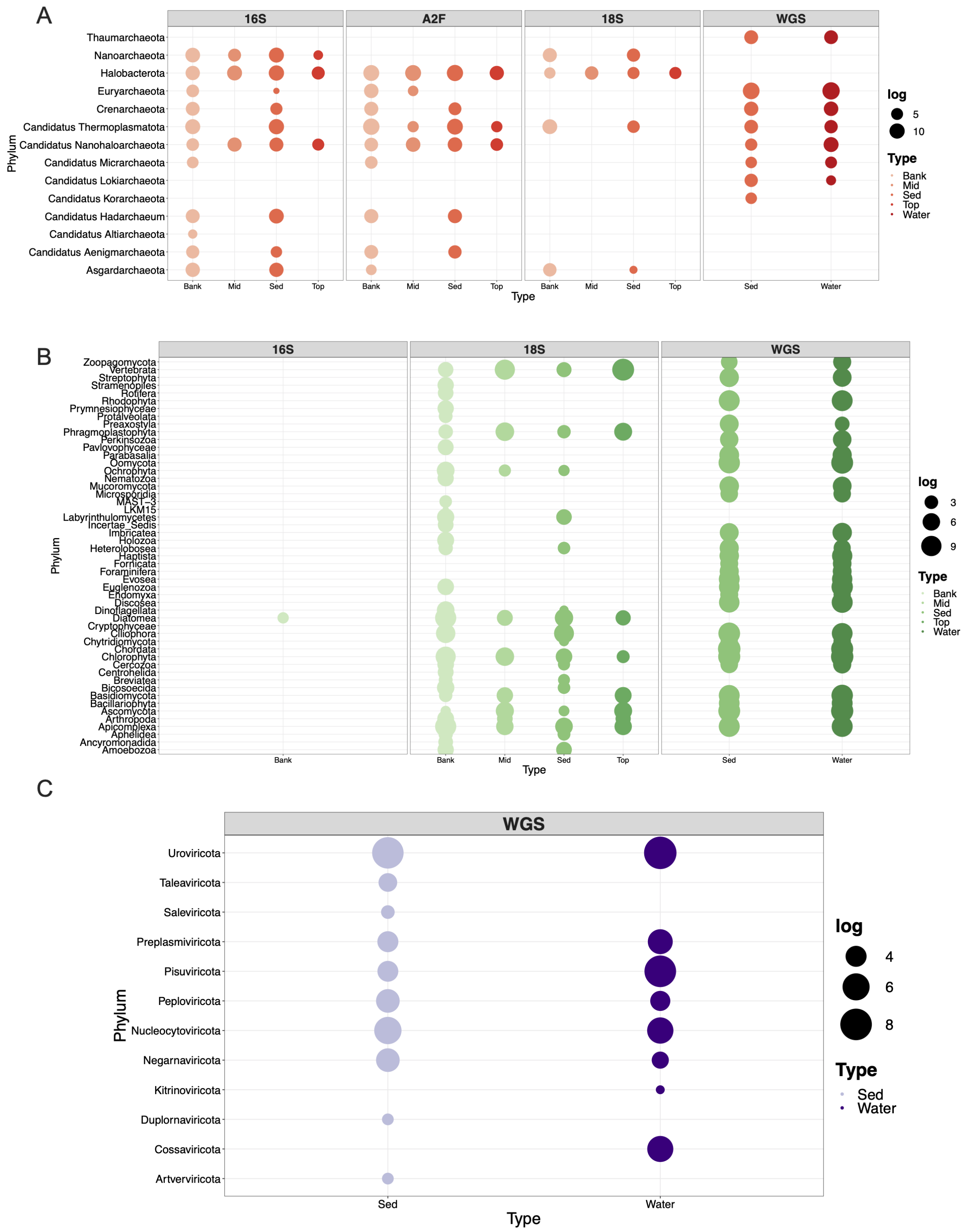


Figure S3: Relative abundance of taxa at the phylum level for Archaea (A), Eukaryota (B) and Viruses (C) from different sample types. Size of dots represents abundance of taxon.

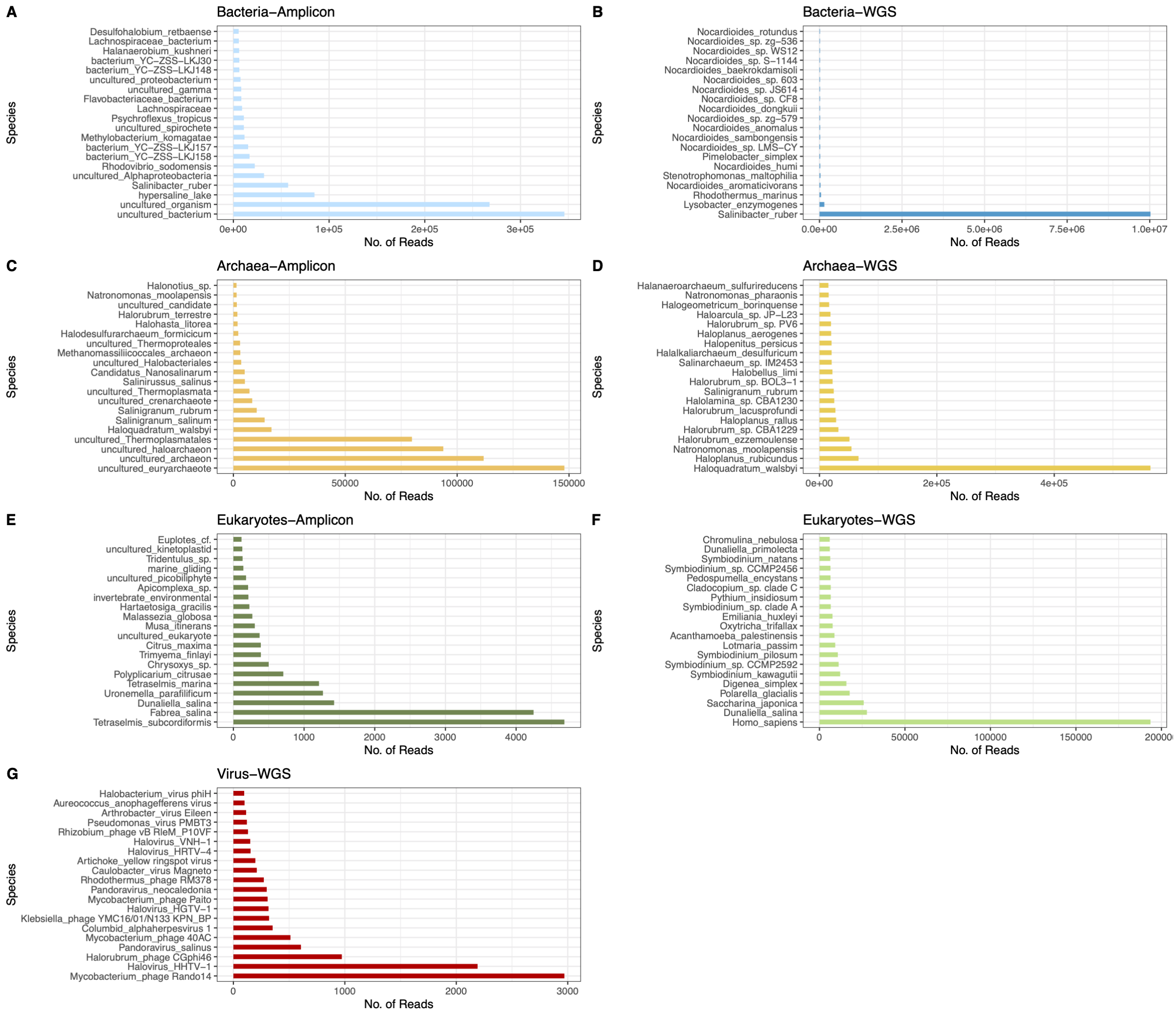


Figure S4: Number of reads of top 20 most abundant species found by amplicon and whole genome sequencing (WGS) sequencing methods in Bacteria (A-B), Archaea(C-D), Eukaryotes(E,F), Virus(G).

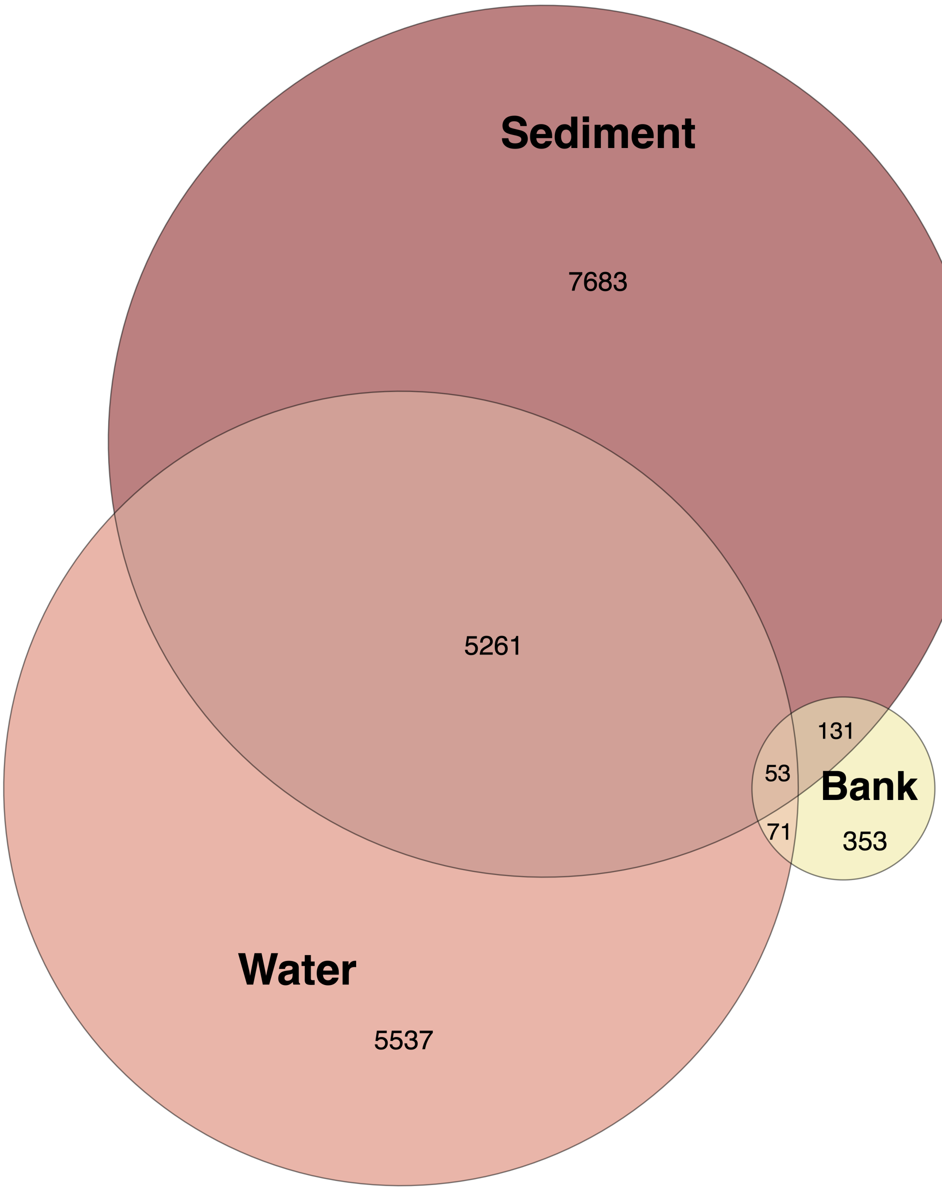


Figure S5: Species overlap between sample types and origin.

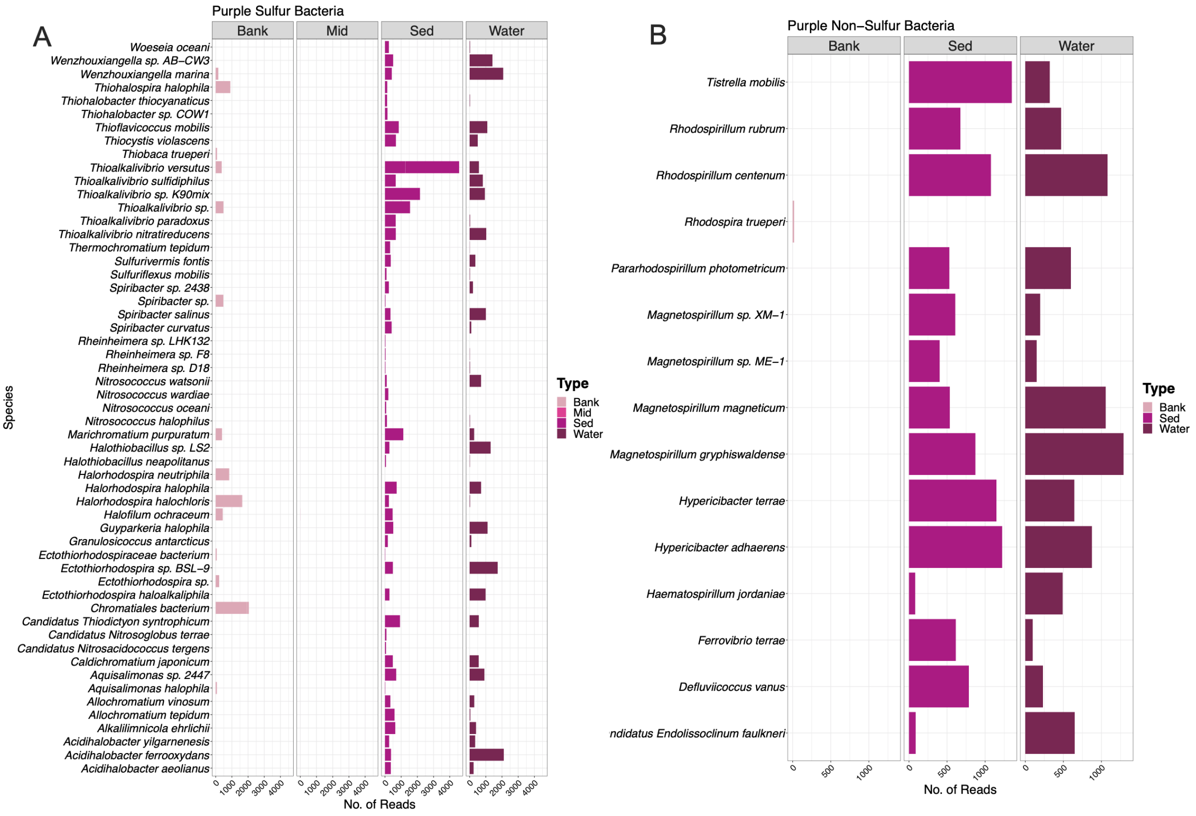


Figure S6: Number of reads of Purple sulfur and non-sulfur bacteria present in Bank, Sediment and Water.

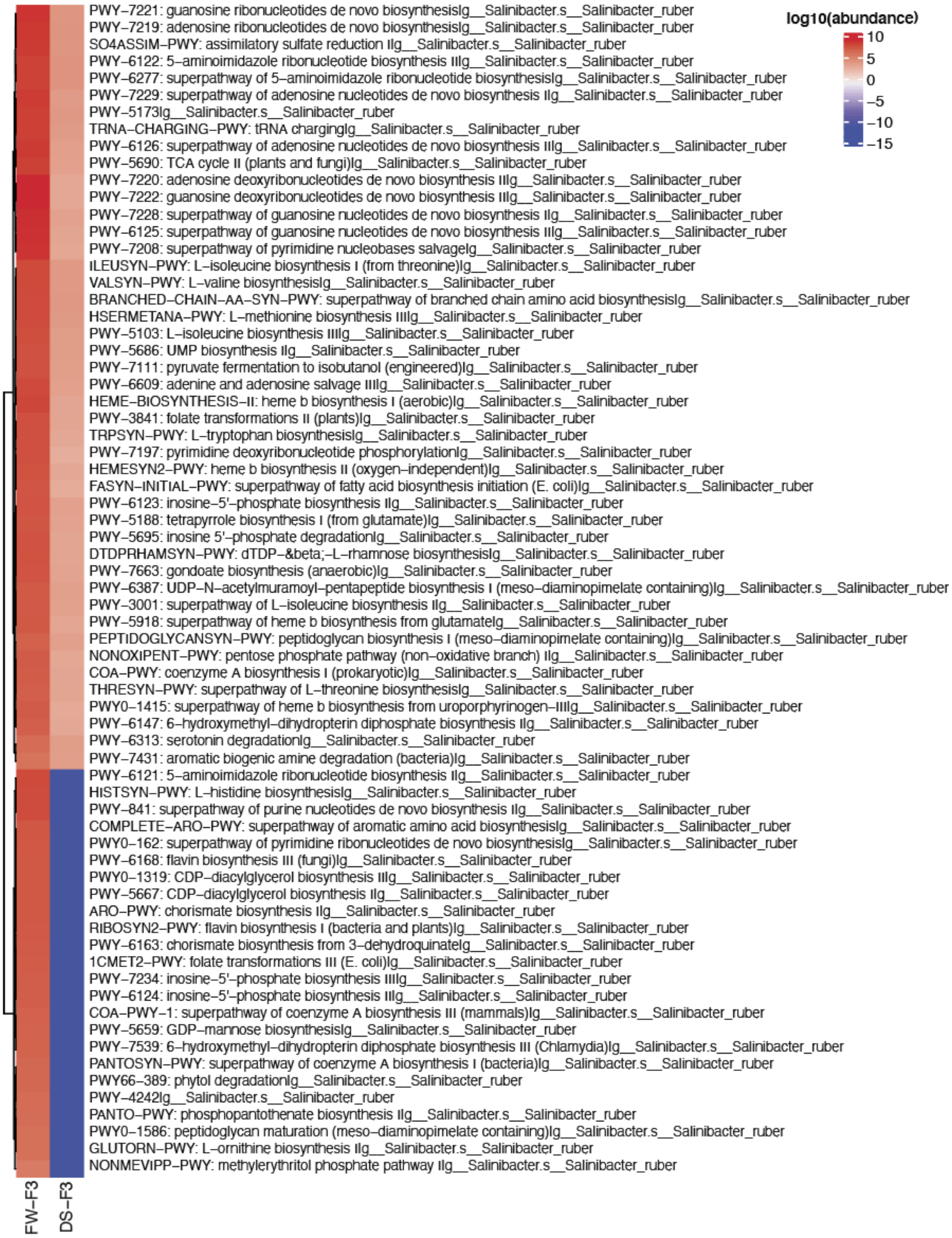


Figure S7: Difference in pathways from the two metagenomes from water (FW) and sediment (DS) in *Salinibater*.

A

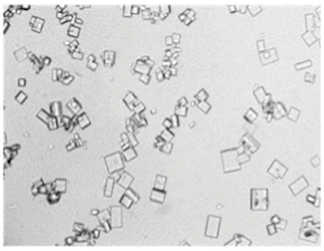

▲ Water sample of Lake Hillier, magnification (200X) reveals the consistency is heavy with salt

B

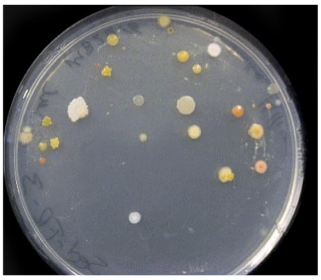

▲ 20uL of sediment sample, plated on Marine Broth Agar 2216

C

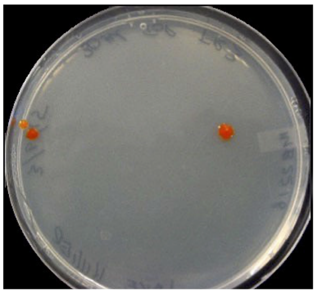

▲ 30uL of water sample, plated on Marine Broth Agar 2216

D

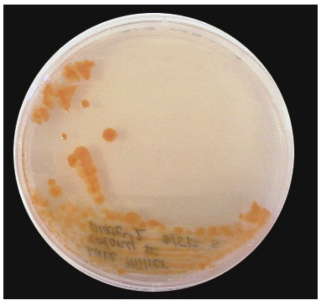

▲ Isolated *Psychroflexus sp,* from water sample culture

E

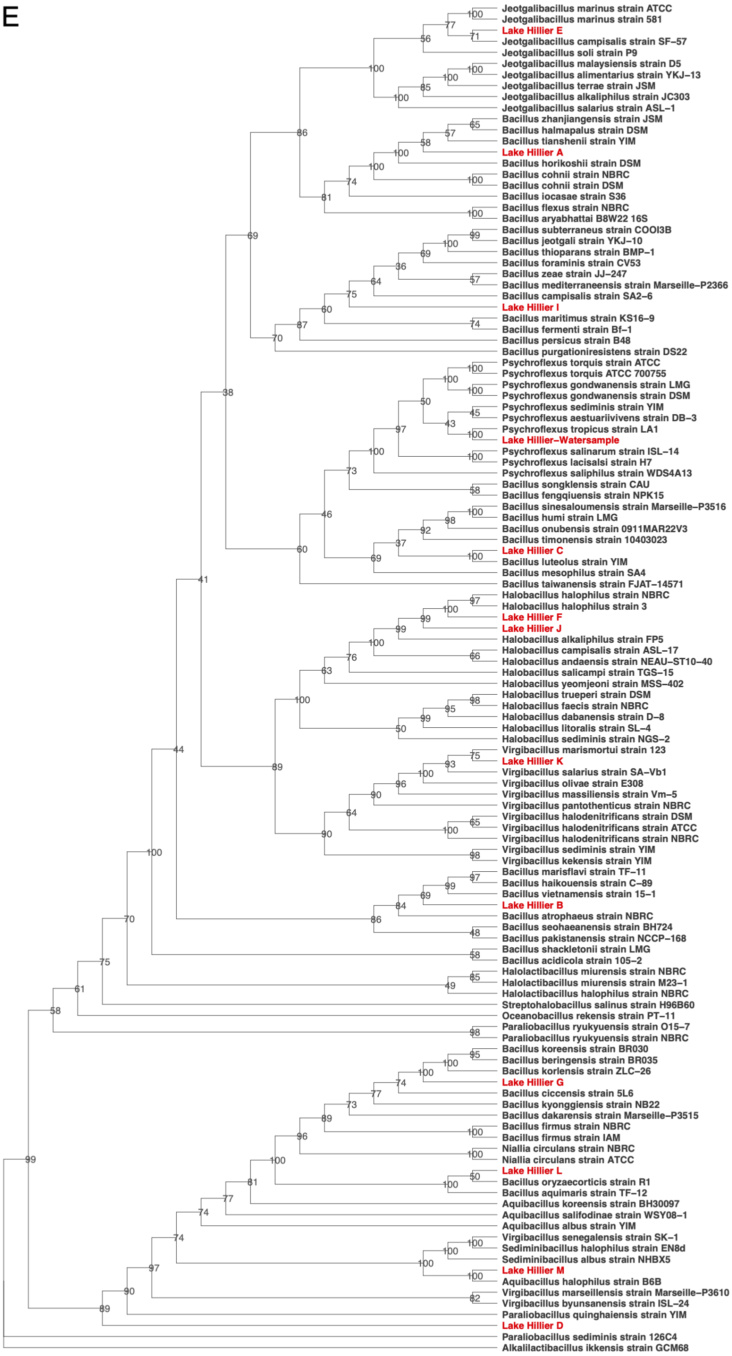


Figure S8: A. Microscopic images of salt crystal from water samples. (B-D) Culturing of Lake Hillier sediment and water samples. E. Maximum Likelihood tree based on 16S gene sequences of isolates (n=13, marked in red) and its closest species.

#### Acknowledgements

The authors would like to thank the support of the Western Australia Department of Parks and Wildlife (DPaW), and John Lizamore and Don Cater for permits and assistance with sampling the lake, as well as the Australian Genome Research Facility (AGRF) and Bioplatforms Australia for infrastructure support. We would like to thank Phil Hugenholtz (University of Queensland) for supporting the expedition to Lake Hillier. Additionally, RNA sequencing was provided by Weill Cornell Medicine, with the support of Jorge Gandara, which was ultimately excluded from the study. We would like to recognize Audria Greenwald (UVM) for their processing and DNA extractions of the water and sediment samples from Lake Hillier. We also acknowledge Jaden J. A. Hastings (WCM) and Deena Najjar (WCM) for their valued input on the project. Additionally, Illumina, Inc. is acknowledged for providing sequencing support for the WGS and RNA-seq data.

#### Funding

The work was supported by Scott Tighe, at the University of Vermont. The expedition to Lake Hillier was funded by the University of Queensland in Brisbane, Australia.

#### Ethical Approval and Consent to participate

"Not applicable"

#### Abbreviations

XMP: Extreme Microbiome Project
AGRF: Australian Genome Research Facility
WGS: Whole Genome Sequencing
TMD: The Microbe Directory
GTDB: Genome Taxonomy Database
PSB: Purple sulfur-bacteria
NPSN: Purple non-sulfur-bacteria
MAGs: Metagenome-Assembled Genomes
ANI: Average Nucleotide Identity
BGCs: Biosynthetic Gene Clusters
GCF: Gene Cluster Family

#### Availability of data and materials

Sequences and scripts will be made available upon publication.

#### Competing interests

B.T.T. consults for Seed Health and Bioscience on microbiome study design and statistical analysis.

#### Consent for publication

All authors read and approved this manuscript.

#### Authors' contributions

The project originated under S.W.T., C.E.M. and K.M. with assistance from M.G.R.G. K.M obtained funding, logistics and permits for sampling. K.M. conducted the field sampling. S.W.T. performed DNA extraction. S.W.T., S.G., W.K.T., and J.R. provided support for library preparation, sequencing and initial analyses. N.J.B conducted 16S rRNA sequencing. D.B. coordinated sample processing and maintaining data storage. K.A.R. provided input for methods and materials. M.A.S.,B.T.T.,J.F., E.A., and C.B. analyzed the data. M.A.S., B.T.T. and K.A.R wrote the manuscript. All authors read and approved the final manuscript.
